# Sortase A-mediated farnesylation of Cdc42 *in vitro*

**DOI:** 10.1101/2024.11.29.626060

**Authors:** Sophie Tschirpke, Nynke M. Hettema, Benjamin Spitzbarth, Rienk Eelkema, Liedewij Laan

## Abstract

Cdc42, a Rho-family GTPase, plays a pivotal role in establishing polarity in *Saccharomyces cerevisiae* by accumulating on the membrane at the site of bud emergence. Cdc42’s ability to bind to membranes, mediated by prenylation, is essential for its function. Prenylation involves either the post-translational addition of a 15-carbon farnesyl group or a 20-carbon geranylgeranyl group to Cdc42’s C-terminus. One of the mayor challenges in studying the biophysical and biochemical interactions of Cdc42 at the polarity spot *in vitro* is obtaining prenylated Cdc42, due to labor-intensive and not easily reproducible traditional methods. Here, we present a streamlined, Sortase A-based approach to farnesylate Cdc42 *in vitro*. This method leverages *E. coli*-expressed Cdc42 with a Sortase A recognition motif, facilitating efficient and accessible farnesylation and purification using a purification tag-based strategy. The farnesylated Cdc42 retains functionality, as evidenced by GTP-dependent membrane binding, making it suitable for further biophysical and biochemical investigations. Additionally, our method can be easily adapted to yield geranyl-geranylated Cdc42.

## Introduction

Small GTPases are highly conserved proteins and are involved in cell polarization in most eukaryotes [Diepeveen et al., 2018]. In *Saccharomyces cerevisiae*, polarity establishment is regulated by the membrane-binding Rho-family GTPase Cdc42, and initiated by its accumulation at the site of bud emergence. Accumulation of Cdc42 on the membrane is driven by at least two interconnected regulatory feedback loops [Chiou et al., 2017, Woods and Lew, 2019, Daalman et al., 2020] and relies on Cdc42’s ability to bind to membranes, which is facilitated by a post-translational modification called prenylation. Here a lipid group is covalentely appended to Cdc42’s C-terminus. Depending on the lipid added (15-carbon farnesyl or 20-carbon geranylgeranyl), prenylation refers to farnesylation or geranyl-geranylation [Caplin et al., 1994, Koch et al., 1997, Coxon and Rogers, 2003, Wang and Casey, 2016].

Over the last decades many *in vivo* and *in vitro* studies have been performed to dissect the molecular interactions that regulate Cdc42’s GTPase activity and Cdc42’s role in polarity establishment [Chiou et al., 2017, Meca et al., 2019, Miller et al., 2020, Chiou et al., 2021, Guan et al., 2023]. Many studies have focussed on regulation of Cdc42 in the cytosol, rather than membrane bound Cdc42. A major challenge for studying membrane-bound Cdc42 *in vitro* is obtaining prenylated Cdc42. Purification of prenylated Cdc42 cannot be achieved using standard *Escherichia coli*-based recombinant expression systems, as prenylation is a eukaryotic post-translational modification for which *E. coli* lacks the required enzymatic machinery. Sofar, prenylated Cdc42 was obtained though (1) insect cell expression systems [Zheng et al., 1994, Zhang and Zheng, 1998, Zheng et al., 1995, Zhang et al., 1999, Kozminski et al., 2003, Johnson et al., 2009, Johnson et al., 2012], (2) purification of membranebound Cdc42 from yeast [Das et al., 2012, Rapali et al., 2017], (3) *in vitro* geranyl-geranylation of recombinantly expressed Cdc42 [Golding et al., 2019], or (4) *E. coli* cell-free prenylated protein synthesis systems [Sonal et al., 2022] (S1 Fig. 1). These approaches are effective but can be difficult to implement: (1) Insect cell expression systems require cell culturing facilities that are not available at every research location. (2) Yeast purification does not allow for Cdc42 over-expression and results in the purification of a mixture of geranyated and farnesylated Cdc42. (3) *in vitro* geranyl-geranylation of Cdc42 requires additional purification and testing of the prenyltransferase geranylgeranyltransferase type-I (GGTase-I), which can vary in activity across different batches. (4) Setting up *E. coli* cell-free prenylated protein synthesis systems demands considerable time, especially for establishing efficient purification techniques for prenylated Cdc42. Further, these methods are not always as accessible and reproducible for researchers from a non-biochemistry group as needed, which hinders scientific progress - despite trying a range of these methods, along with other novel approaches, we did not achieve success with any of them (summarized in S1).

Here, we utilize the transpeptidase Sortase A (which we will refer to as ‘Sortase’ in the following) from *Staphylococcus aureus* to prenylate Cdc42 in a straightforward *in vitro* reaction (Fig. 1). Sortase is a versatile enzyme used for protein labeling due to its ability to perform both cleavage and ligation in one step: It recognizes the LPXTG motif (where L is leucine, P is proline, X is a variable amino acid, T is threonine, and G is glycine) in a substrate, cleaves the bond between threonine and glycine in this motif, and subsequently catalyzes the formation of an amide bond between the threonine of the substrate and the *N*-terminal glycine of an oligoglycine probe. Sortase has almost no requirements for the structure or sequence of the probe (as long as it contains oligoglycine), and even accepts a wide range of non-proteinaceous molecules, like fluorescent dyes, fatty acids, nucleic acids, polymers, or C14-C24 lipid chains [Pritz et al., 2007, Antos et al., 2008, Antos et al., 2017, Popp and Ploegh, 2011]. We utilized the versatility of Sortase to produce one type of prenylated Cdc42 - farnesylated Cdc42. In the same way, our method can be used to yield the other type of prenylation: geranyl-geranyltion. To do so, we purified recombinantly expressed Cdc42, in which a Sortase recognition motif is inserted in-between Cdc42’s C-terminus and a *C*-terminal purification tag (Fig. 1), and synthesized farnesylated triglycine (Fig. 4). Such peptides were not commercially available at that time, but several companies offer them now. We then performed a Sortase-mediated *in vitro* reaction, in which Cdc42’s *C*-terminal purification tag is replaced by the farnesylated triglycine (Fig. 1). Our approach offers several key advantages: (1) It utilizes recombinant Cdc42 expressed in *E. coli*, eliminating the need for specialized cell culturing skills and facilities. (2) It employs the commercially available Sortase enzyme, avoiding the need for additional enzyme purification. (3) It leverages a tag-based purification strategy for the straightforward purification of prenylated Cdc42. Conversely, a drawback of this technique is that it incorporates a small linker, composed of parts of the Sortase recognition motif and the oligoglycine from the probe, between Cdc42’s *C*-terminus and the farnesyl group (Fig. 1).

**Figure 1.**
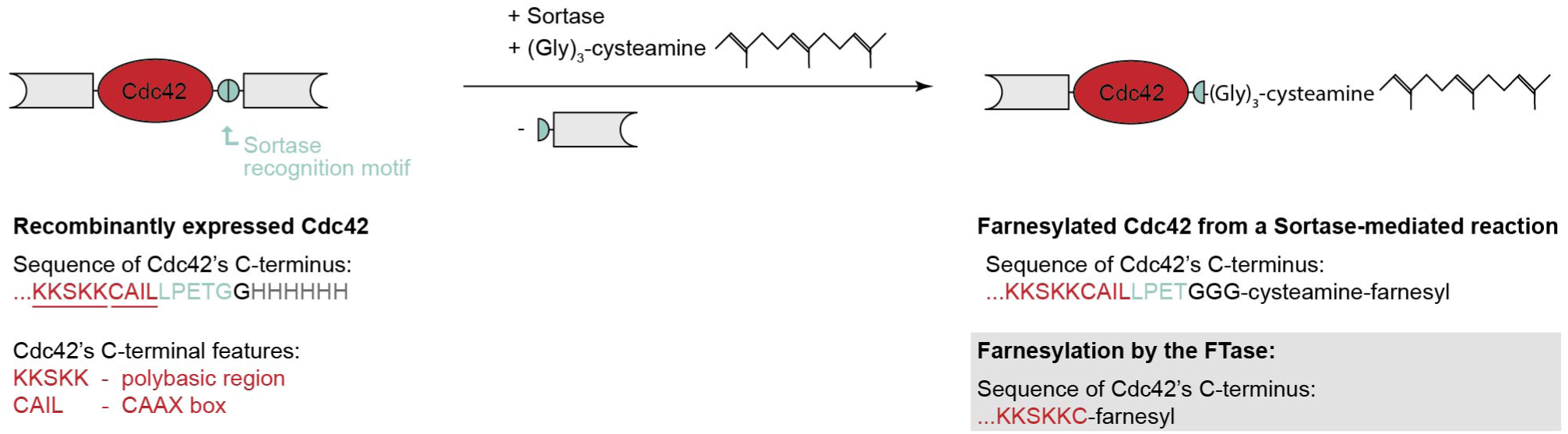
Schematic illustration of Sortase-mediated farnesylation of recombinantly expressed Cdc42. The sequence of Cdc42’s *C*-terminus, before and after Sortase-mediated farnesylation, is shown below the illustration. *C*-terminal Cdc42 features are annotated in red and the Sortase recognition motif is highlighted in green. For comparison, the sequence of Cdc42’s *C*-terminus after farnesylation with farnesyltransferase (FTase), as occurring *in vivo*, is given.

Our data shows that Sortase-mediated reactions are an effective method for farnesylating Cdc42, which can be easily be purified through a purification-tag-based approach. The addition of the detergent Brij58 aids the reaction yield, presumably through solubilizing farnesylated Cdc42. Further, farnesylated Cdc42 binds to membranes in a GTP-dependent manner, suggesting that the linker region between the protein’s *C*-terminus and the farnesyl group does not impede protein functionality.

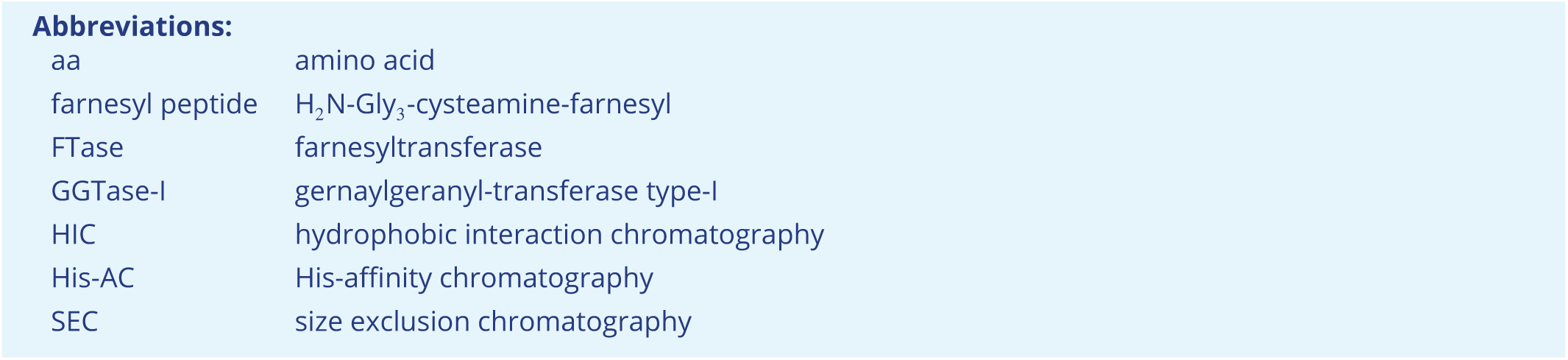

## Results

### Cdc42 can easily be farnesylated in a Sortase-mediated reaction

We purified Cdc42 with a *C*-terminal Sortase recognition motif from *E. coli* and synthesized H_2_N-Gly_3_-cysteamine-farnesyl (’farnesylated triglycine’, Fig. 4) ^1^, and used a His-tagged Sortase mutant with improved catalytic properties ([Chen et al., 2016], available from BPS Bioscience) for the labeling reactions (Fig. 2a).

**Figure 2.**
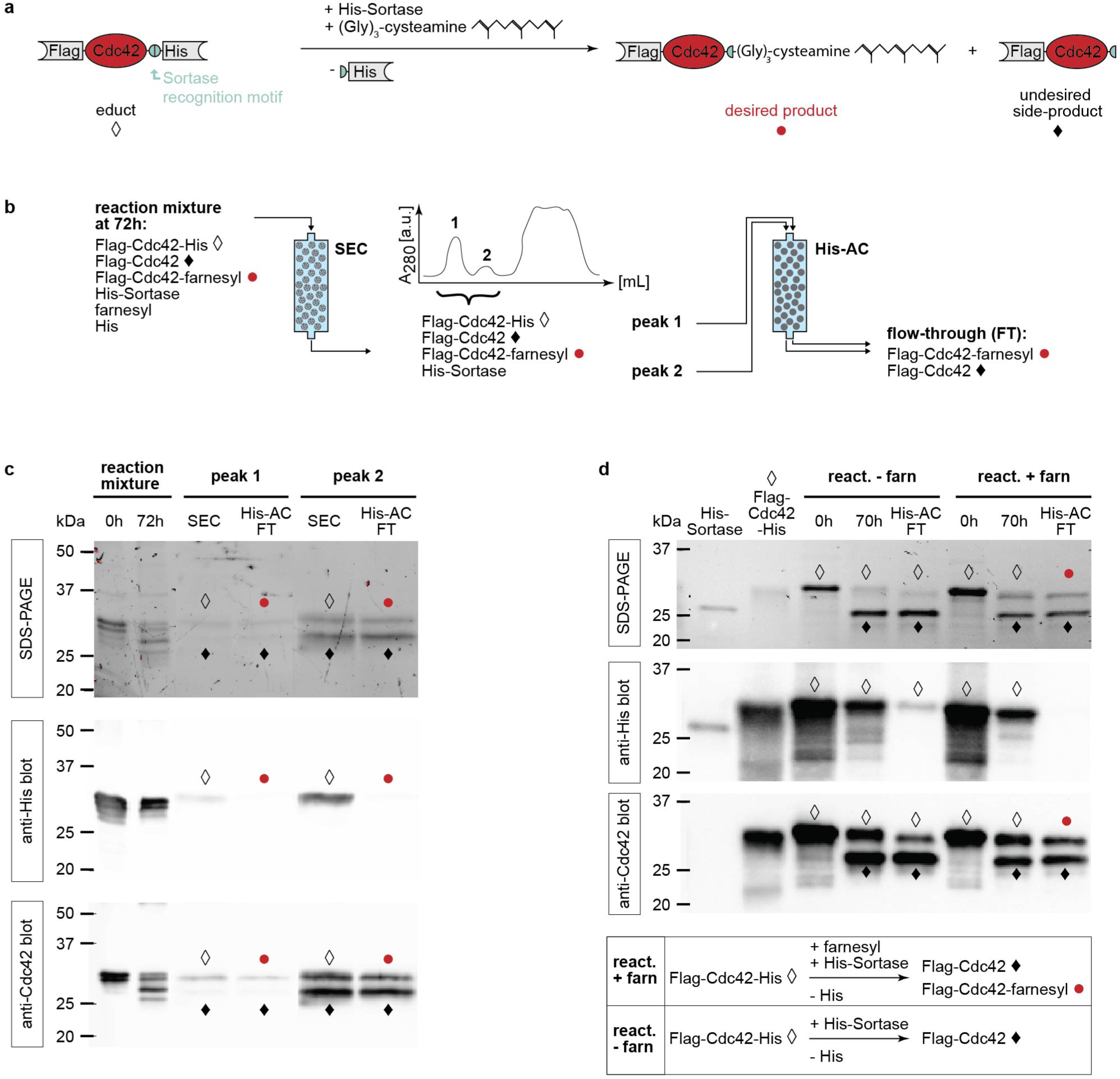
Cdc42 can be farnesylated in a Sortase-mediated reaction. (a) Schematic illustration of the Sortase-mediated labeling reaction of double-tagged Cdc42 with farnesylated triglycine. (b) Clean-up procedure for the labeling reaction: after an incubation of 72 h size exclusion chromatography (SEC) is used to separate unreacted farnesylated triglycine and cleaved purification His-tags from proteins, which elute as two peaks (peak 1 and 2). Each SEC peak is then loaded onto a His-affinity chromatography (His-AC) column. Labelled protein (Flag-Cdc42-farnesyl) and reacted but unlabelled protein (Flag-Cdc42) do not bind to the column due to the absence of a His-tag. (c) SDS-PAGE and Western blot analysis of the reaction mixture pre-reaction (0 h), post-reaction (72 h), of the SEC peaks and the His-AC flow-through: Both SEC peaks contain all three Cdc42 species (Flag-Cdc42-His, Flag-Cdc42-farnesyl, Flag-Cdc42). After His-AC a mixture of Flag-Cdc42-farnesyl and Flag-Cdc42 remains. (d) SDS-PAGE and Western blot analysis of a Sortase-mediated reaction with and without farnesylated triglycine that was only cleaned-up by His-AC. The control reaction absent of farnesylated triglycine confirms that the ∼28 kDa band is Flag-Cdc42.

To optimize reaction conditions, we carried out screens using Bem1 and Cdc42 and Alexa488 peptide (S2). Initially we tried monitoring product formation of Cdc42-farnesyl using anti-farnesyl Western blotting, but an eight condition screen utilizing materials stated in [Kennedy et al., 2019, Li et al., 2020] revealed that all conditions gave false-positive and false-negative signal (data not shown). Considering the wide use and probe tolerances of Sortase, we assumed that the general trends of Cdc42/Bem1 Alexa488 peptide condition screens are applicable to reactions of Cdc42 and farnesylated triglycine. We monitored the formation of labeled protein through the Alexa488-signal of the protein bands on SDS-PAGE (S2). The more intense the signal, the more protein got labeled. Cdc42 and Bem1 Alexa488 peptide screens showed the same trends: Increasing amounts of Sortase enzyme, peptide probe, and reaction time lead to an increased signal, and thus increased product formation. Reactions carried out for 72 h at 4^◦^C at a 2:100:2000 molar ratio of Sortase : protein : Alexa488 peptide lead to the highest amount of product formation. For the reaction of Cdc42 with farnesylated triglycine we used this condition, and 5× higher peptide concentrations to further boost product formation. We considered even longer incubation times as they might increase the yield, but decided against it as this also increases the chances for protein degradation and aggregation. We then used the Sortase-mediated reaction to ligate farnesylated triglycine to double-tagged Cdc42 (Flag-Cdc42-His, Fig. 2a) and utilized the presence/absence of the His-tag for cleaning-up the reaction mixture (Fig. 2b). During the labeling reaction the Sortase enzyme cleaves Cdc42’s *C*-terminal His-tag and ligates farnesylated triglycine to the protein. The final reaction mixture therefore contains three Cdc42 species:

- educt: Flag-Cdc42-His, 29 kDa (marked with a diamond in Fig. 2)
- desired product: Flag-Cdc42-farnesyl, 28.5 kDa (marked with a red dot in Fig. 2)
- undesired side-product: Flag-Cdc42, 28 kDa (marked with a filled diamond in Fig. 2)

Additionally, Sortase (22 kDa, migrating at 26 kDa on SDS-PAGE (Fig. 2d)), cleaved *C*-terminal His-tags, and remaining farnesylated triglycine are part of the reaction mixture (Fig. 2b).

As a first step, size exclusion chromatography (SEC) was used to separate reactants and products by size. Considering the proteins’ sizes, SEC separates a mixture of the three Cdc42 species and Sortase enzyme from remaining farnesylated triglycine and cleaved His-tag peptides. Interestingly, in the presence of the detergent Brij58, two such peaks are observed (Fig. 2b), both containing all three Cdc42 species (Fig. 2c). The appearance of two peaks is detergent-dependent; in the absence of detergent, only peak 2 is present, while using CHAPS results in peak 2 with a broad shoulder in the region where peak 1 would normally appear (S3). Next, the two SEC peak fractions were loaded repeatedly on a nickel column and the flow-through was collected (His-affinity chromatography (His-AC)). His-Sortase and Flag-Cdc42-His bind to the column material, leaving only the two reaction products (Flag-Cdc42-farnesyl, Flag-Cdc42) in the flow-through (Fig. 2b).

To qualify and quantify the labeling success, we analyzed all reaction and purification steps by SDS-PAGE and anti-His and anti-Cdc42 Western blotting (Fig. 2c): Bands that are visible on both anti-His and anti-Cdc42 blots correspond to the educt (Flag-Cdc42-His, diamond). Bands that are visible on the anti-Cdc42 blot, but not on the anti-His blot, correspond to either the undesired side-product (Flag-Cdc42, filled diamond) or farnesylated Cdc42 (red dot). After 72 hours of reaction, four distinct bands were observed on SDS-PAGE (Fig. 2c): a double band at 30 and 29.5 kDa, along with bands at 27 and 26 kDa. The double band was also present in the 0 hour sample (taken at the start of the reaction) and appeared on both the anti-Cdc42 and anti-His blots, identifying it as the educt (Flag-Cdc42-His). The 27 and 26 kDa bands were visible on the anti-Cdc42 blot but not the anti-His blot, suggesting they represent the reaction products (Flag-Cdc42 and Flag-Cdc42-farnesyl). Both SEC peaks showed two bands - ∼30 and ∼28 kDa - likely because some of the bands merged during this step (Fig. 2c). A change in buffer conditions could explain why these bands migrated at a slightly higher molecular weight than before. Interestingly, the signal for SEC peak 1 was significantly weaker than that of peak 2 on SDS-PAGE, despite peak 1 having a higher absorbance during SEC. As in the 72 hour sample, the 30 kDa band from SEC peak 1 was visible on both anti-His and anti-Cdc42 blots, identifying it as the educt Flag-Cdc42-His. The 28 kDa band, detectable only on the anti-Cdc42 blot, likely corresponds to one of the reaction products. We performed a control reaction without the farnesylated triglycine, which results in a mixture of Flag-Cdc42-His (educt) and Flag-Cdc42 (side-product) (Fig. 2d). This reaction was purified using only His-AC. The SDS-PAGE analysis showed a weak 30 kDa band and an intense 28 kDa band. The 30 kDa band was visible on the anti-His blot, while the 28 kDa band was not, indicating that the 28 kDa band corresponds to the side-product Flag-Cdc42. ^2^ After His-AC, the flow-through should contain only Flag-Cdc42 and Flag-Cdc42-farnesyl. The band patterns of both SEC peaks in the His-AC flow-through again showed the 30 and 28 kDa bands (Fig. 2c). Notably, neither band was visible on the anti-His blot, indicating both are His-tag-free. Since the control reaction identified the lower band as Flag-Cdc42, the 30 kDa band must correspond to the desired product, Flag-Cdc42-farnesyl, which migrates similarly to the educt. We prepared His-AC flow-through samples for mass analysis (Max Perutz Labs, Vienna), which confirmed the presence of Flag-Cdc42-farnesyl in both sec peaks (S4), confirming our Western blot analysis.

Starting with 3.4 mg of Cdc42 for the reaction, we found that the His-AC flow-through from SEC peak 1 contained 1.7 mg of protein (50% of total Cdc42), and SEC peak 2 contained 0.8 mg (24% of total Cdc42), which matches the SEC absorbance data. Both peaks contain a mixture of Flag-Cdc42 and Flag-Cdc42-farnesyl. In SEC peak 1, the ratio of the 30 kDa Flag-Cdc42-farnesyl band to the 28 kDa Flag-Cdc42 band was approximately 2:1, while in SEC peak 2 it was around 1:2. This indicates that 0.85 mg (33% of total Cdc42) of farnesylated Cdc42 was present in peak 1, and 0.27 mg (8% of total Cdc42) in peak 2, leading to a total labeling efficiency (after all clean-up steps) of roughly 41%.

Our findings indicate that SEC peak 1 contains most of the farnesylated protein, with the peak only appearing when Brij58 is present. This is in agreement with literature, showing that Brij58 increases the solubility of prenylated Cdc42 [Sonal et al., 2022]. The weaker SDS-PAGE signal for SEC peak 1 may be due to farnesylated Cdc42 adhering to the surfaces of tubes and pipettes, leading to loss during handling. This implies that the actual labeling efficiency before clean-up steps is higher than 41%. Our purification strategy yields two samples; one with a 2:1 ratio of Flag-Cdc42-farnesyl to Flag-Cdc42 (SEC peak 1), and another with the reverse ratio (SEC peak 2). We also explored a hydrophobicity-based purification approach, but were unable to effectively separate farnesylated from not farnesylated Cdc42 (S5).

Taken together, our data demonstrate that Sortase-mediated reactions provide an efficient method for farnesylating Cdc42. The presence of affinity tags facilitates purification, and the addition of the detergent Brij58 significantly boosts labeling efficiency, likely by improving solubility of the farnesylated Cdc42.

### Membrane-binding of farnesylated Cdc42

To check whether the prenylated Cdc42 had functional membrane binding, we carried out a liposome co-floatation assay. Here, liposome-bound protein is distinguished from the unbound protein by centrifugation through a sucrose gradient. We mixed Cdc42 that previously underwent a Sortase-mediated reaction with or without the farnesylated triglycine peptide with negatively charged liposomes (80% phosphatidylcholine, 20% phosphatidylserine) into the first layer of high sucrose density and added two steps of lower densities on top. An excess of GTP was added to ensure an active state of Cdc42. After centrifugation at high speed, the top and bottom fraction were analysed through an anti-Cdc42 blot (Fig. 3a). We found that there is almost no protein present in fraction A on an anti-Cdc42 blot for the control reaction product, Sortase-mediated reaction without farnesylated triglycine peptide (Fig. 3b). In fraction B, for the control reaction product, we see a mixture of reaction products Flag-Cdc42 and Flag-Cdc42-His as seen in (Fig. 2d). The absence of the control reaction product in the top fraction, indicates that no protein is bound to the liposomes, suggesting that without prenylation, Cdc42 does not bind membranes. For the Flag-Cdc42-farnesyl sample, on an anti-Cdc42 blot, we see the majority of protein in fraction A, indicative of protein-membrane binding (Fig. 3b). The near absence of Flag-Cdc42-farnesyl in the bottom fraction suggest that the protein binds the membrane strongly and is rarely present in unbound form under the influence of GTP.

**Figure 3.**
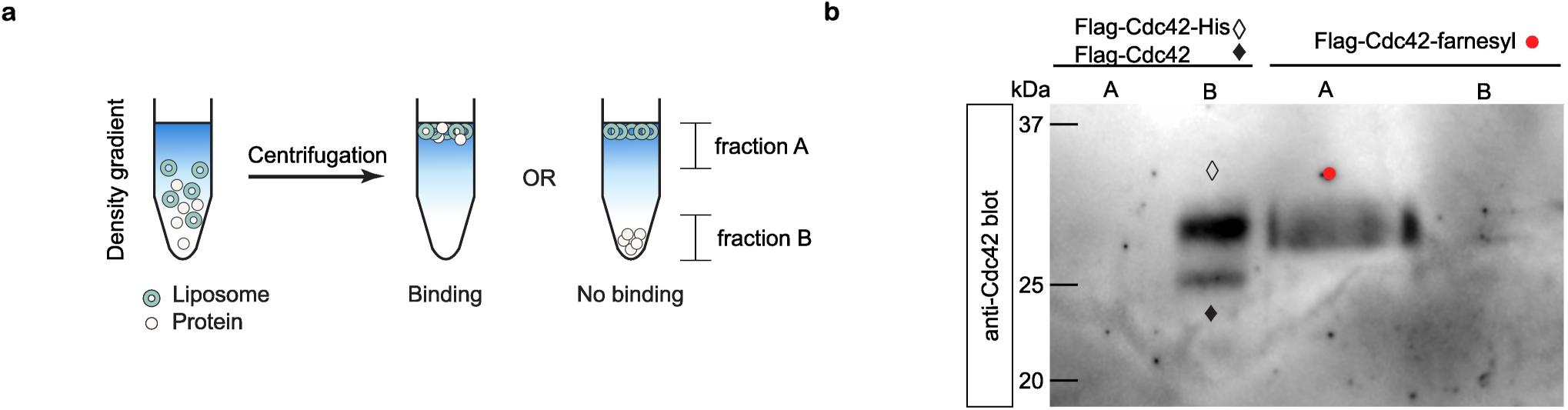
Farnesylated Cdc42 can bind membranes in liposome co-floatation assay. (a) Schematic illustration of the liposome co-floatation assay. (b) Western blot analysis of the top and bottom fraction of co-floatation assay conducted with protein from a Sortase-mediated reaction with and without farnesylated triglycine, cleaned by His-AC. The control reaction absent of farnesylated triglycine is visible as two bands on the blot; a lower band with successful Sortase cutting (Flag-Cdc42) and a higher band with unsuccessful Sortase cutting (Flag-Cdc42-His).

## Discussion

In this study, we utilized a Sortase-mediated reaction to farnesylate Cdc42. One of the major challenges we encountered was the detection of the farnesylated product, as previously used Western blotting protocols proved ineffective [Kennedy et al., 2019, Li et al., 2020]. To address this, we adopted an indirect approach by blotting for all Cdc42 species, including the educt, and used a control reaction to identify the undesired side-product. This allowed us to exclude bands corresponding to nonfarnesylated Cdc42 species and accurately identify the farnesylated product on the gel/blot (Fig. 2). A more straightforward approach would be to implement an anti-farnesyl blotting protocol, which, if reliably developed, could significantly benefit the wider scientific community. Alternatively, using a fluorescently labeled farnesylated triglycine, similar to the approach by Sonal *et al*. [Sonal et al., 2022], would allow to directly detect and measure the formation of the farnesylated Cdc42. Due to the limitations of blotting, we verified the presence of farnesylated Cdc42 using mass-spectrometry. However, only the intact protein mass-spectrometry analysis was successful. Earlier attempts to detect a farnesylated peptide through a peptide mapping approach (i.e., digesting Cdc42 with GluC) yielded unreliable data, likely due to peptide loss during processing and uneven digestion, as GluC appeared to have preferred cleavage sites (S1). In summary, detecting farnesylation remains difficult, likely due to the opposing characteristics of the strongly hydrophobic farnesyl group and the comparatively hydrophilic protein, and more reliable methods are needed.

In addition to the challenges of detecting farnesylated Cdc42, working with it without significant loss is difficult. We suspect that a substantial portion of the farnesylated Cdc42 is lost during the labeling and purification processes. This is suggested by the observation that, despite high absorbance during SEC, the gel of the SEC peak 1 (containing the majority of farnesylated Cdc42) shows only a faint signal (Fig. 2). We found that using low-binding materials (such as tubes and pipette tips) helped reduce the overall loss of farnesylated Cdc42. Given these challenges, we believe that an overall farnesylation yield of ∼40% after all purification steps is high.

In addition to the Sortase-mediated reaction, we also attempted to express Cdc42 in insect cells, yeast, and *E. coli* coexpressing farnesyltransferase (FTase) and Cdc42 (S1). While expression and farnesylation were successful in all systems, the farnesylated Cdc42 remained stuck in the membrane fraction, and various solubilization methods we tried were unsuccessful (S1). The development of a robust solubilization method would be a worthwhile research endeavor, as it would address the main challenge in these three systems and enhance their usability. A common strategy to prevent prenylated GTPases from being sequestered in the membrane fraction is co-expressing them with guanine nucleotide dissociation inhibitors (GDIs)[Bezeljak et al., 2020]. This, however, introduces the GDI into the system, which may not always be desirable. Given the challenges with *in vivo* expression systems, *in vitro* prenylation appears to be a more viable option. Traditionally, this is achieved using purified FTase and geranylgeranyltransferase-I (GGTase-I) [Golding et al., 2019, Kuhm et al., 2023]. We chose not to pursue this method because it would require purifying additional enzymes and verifying their activity, which, combined with the difficulty of detecting farnesylation, would introduce additional uncertainty.

Instead, we opted for the Sortase enzyme, which is versatile, commercially available, and easily purified (a process supported by well-established purification protocols)[Pritz et al., 2007, Antos et al., 2008, Antos et al., 2017, Popp and Ploegh, 2011]. The drawback of our method is the insertion of a linker region between Cdc42’s *C*-terminus and the farnesyl group, consisting of the Sortase recognition motif and triglycine (Fig. 1). With FTase-mediated prenylation, no linker is introduced: Prenylation targets a conserved *C*-terminal CAAX box, where ‘C’ is a cysteine that undergoes modification, ‘A’ represents aliphatic amino acids, and ‘X’ is any amino acid. A thioether bond forms between the cysteine and a hydrophobic isoprenoid lipid, after which the remaining three amino acids are cleaved, and the cysteine is carboxymethylated[Caplin et al., 1994, Coxon and Rogers, 2003].

Thus, in this process, the farnesyl group is added directly to the cysteine. In contrast, our approach appends the Sortase recognition motif after the CAAX box, resulting in a linker region of ten amino acids (Fig. 1). This linker could be shortened to seven amino acids by reducing the CAAX box to just the cysteine and appending the Sortase motif directly afterward. Nonetheless, we found no indication that the linker impaired Cdc42 functionality: Our earlier investigations demonstrated that neither *C*-terminal tags nor the presence of longer unstructured regions affected Cdc42 GTPase activity or its interaction with the GEF Cdc24 [Tschirpke et al., 2023]. Here, we further show that the linker does not impede membrane binding, indicating that Cdc42 remains functional despite the modification.

This was supported further by the liposome co-floatation assay, where we observed GTP dependent membrane binding of the farnesylated Cdc42. The observed binding of this assay is comparable to the strong Cdc42 binding observed in recent in vitro literature ([Sonal et al., 2022]). However, this assay does not yet show the full binding dynamics of Cdc42. Addition of a GDI into the system could help to show the full cyclical behaviour of binding and unbinding of Cdc42([Johnson et al., 2009, Sonal et al., 2022]). Membrane composition can have a large influence on binding dynamics and was in this case chosen to be comparable to the most recent report on Cdc42 membrane binding (Sonal). Although other reports([Johnson et al., 2009, Das et al., 2012]) show the essence of other negatively charged lipids for reliable Cdc42 membrane binding, we have observed strong GTP dependent binding with a simple two component artificial membrane.

## Materials and Methods

### Plasmid construction

The gene of interest (Cdc42) was obtained from the genome of *Saccharomyces cerevisiae* W303 and was amplified through PCR. The target vector was also amplified through PCR. Additionally, each PCR incorporated a small homologous sequences needed for Gibson assembly [Gibson et al., 2009]. After Gibson assembly, the resulting mixture was used to transform chemically competent Dh5*α* and BL21 DE::3 pLysS cells and plated out onto a Petri dish containing Lysogeny broth agar and the correct antibiotic marker. Construction routes of plasmids and primers used for each PCR, as well as the amino acid sequences of the proteins used in this publication, are stated in S6.

### Buffer composition

If not mentioned otherwise, buffers are of the composition stated in Tab. 2.

**Table 1.** List of protein constructs/plasmids used throughout this publication.

| Plasmid | Description | Source |
| --- | --- | --- |
| pET28a-His-mcm10-Sortase-Flag | template for pRV007 | received from N. Dekker (TU Delft), based on pBP6 [Douglas and Diffley, 2016] |
| pRV007 | His-Cdc42-Flag | this work |
| pRV073 | Flag-Cdc42-His | this work |

**Table 2.**
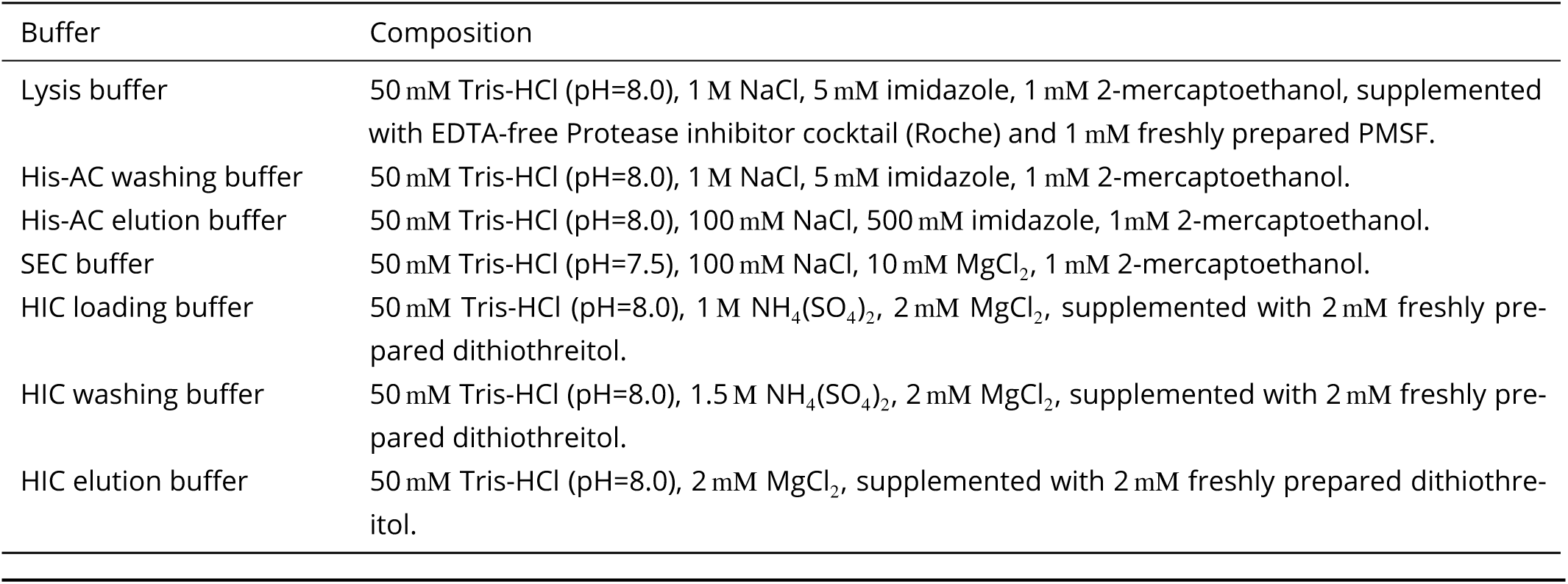
Buffer composition.

| Buffer | Composition |
| --- | --- |
| Lysis buffer | 50 mM Tris-HCl (pH=8.0), 1 M NaCl, 5 mM imidazole, 1 mM 2-mercaptoethanol, supplemented with EDTA-free Protease inhibitor cocktail (Roche) and 1 mM freshly prepared PMSF. |
| His-AC washing buffer | 50 mM Tris-HCl (pH=8.0), 1 M NaCl, 5 mM imidazole, 1 mM 2-mercaptoethanol. |
| His-AC elution buffer | 50 mM Tris-HCl (pH=8.0), 100 mM NaCl, 500 mM imidazole, 1 mM 2-mercaptoethanol. |
| SEC buffer | 50 mM Tris-HCl (pH=7.5), 100 mM NaCl, 10 mM MgCl<sub>2</sub>, 1 mM 2-mercaptoethanol. |
| HIC loading buffer | 50 mM Tris-HCl (pH=8.0), 1 M NH<sub>4</sub>(SO<sub>4</sub>)<sub>2</sub>, 2 mM MgCl<sub>2</sub>, supplemented with 2 mM freshly prepared dithiothreitol. |
| HIC washing buffer | 50 mM Tris-HCl (pH=8.0), 1.5 M NH<sub>4</sub>(SO<sub>4</sub>)<sub>2</sub>, 2 mM MgCl<sub>2</sub>, supplemented with 2 mM freshly prepared dithiothreitol. |
| HIC elution buffer | 50 mM Tris-HCl (pH=8.0), 2 mM MgCl<sub>2</sub>, supplemented with 2 mM freshly prepared dithiothreitol. |

### Protein expression and purification

**Cdc42 (pRV007 and pRV073)** was expressed in Bl21::DE3 pLysS cells. Cells were grown in Lysogeny broth at 37^◦^C until an OD_600_ of 0.7. The expression was induced through addition of 1.0 mM IPTG, after which cells were grown for 3 h at 37^◦^C. Cells were harvested through centrifugation. Cell pellets were resuspended in lysis buffer and lysed with a high pressure homogenizer (French press cell disruptor, CF1 series Constant Systems) at 4^◦^C, using 5-10 rounds of exposing the sample to pressurisation. The cell lysate was centrifuged at 37000× g for 30 min and the supernatant was loaded onto a HisTrap^TM^ excel column (Cytiva). After several rounds of washing with His-AC washing buffer, the protein was eluted in a gradient of His-AC washing buffer and His-AC elution buffer. The protein as dialysed twice in SEC buffer. After the addition of 10% glycerol, samples were flash frozen in liquid nitrogen and kept at -80^◦^C for storage. The proteins are shown on SDS-PAGE in S6.

#### Note on His-AC

HisTrap^TM^ excel column (Cytiva) columns bought 2020 or later required a higher amount of imidazole in the lysis and washing buffer that stated in Tab. 2, as indicated by the recommendation ‘use 20-40 mM imidazole in sample and binding buffer for highest purity’ on the column package. For these columns the amount of imidazole in lysis and His-AC washing buffer was increased to 50 mM.

### Synthesis of H_2_N-Gly_3_-cysteamine-farnesyl (’farnesylated triglycine’)

#### Abbreviations

Boc: *tert*-butoxycarbonyl protecting group
Gly: glycine
DCM: dichloromethane
DMF: *N,N*-dimethylformamide
PyBOP: benzotriazol-1-yl-oxytripyrrolidinophosphonium hexafluorophosphate
DiPEA: *N,N*-diisopropylethylamine
TFA: trifluoroacetic acid
MeOH: methanol
NMR: nuclear magnetic resonance
DMSO: dimethylsulfoxide

#### General Information

Boc-Gly_3_-OH was purchased from Bachem. DCM, DMF, PyBOP, DiPEA, Cysteamine hydrochloride, farnesyl bromide, TFA, 7 M NH_3_ in MeOH, MeOH were purchased from Sigma Aldrich. DMSO-*d*_6_ was purchased from Euriso-top. Deionised (milliQ) water was made in our laboratory. Unless stated otherwise, all chemicals were used as received. For all synthetical steps anhydrous solvents were used. NMR spectra were recorded on an Agilent-400 MR DD2 (399.67 MHz) instrument, NMR measurements were taken at 298 K.

#### Synthesis

An overview of the synthesis steps is given in Fig. 4.

**Figure 4.**
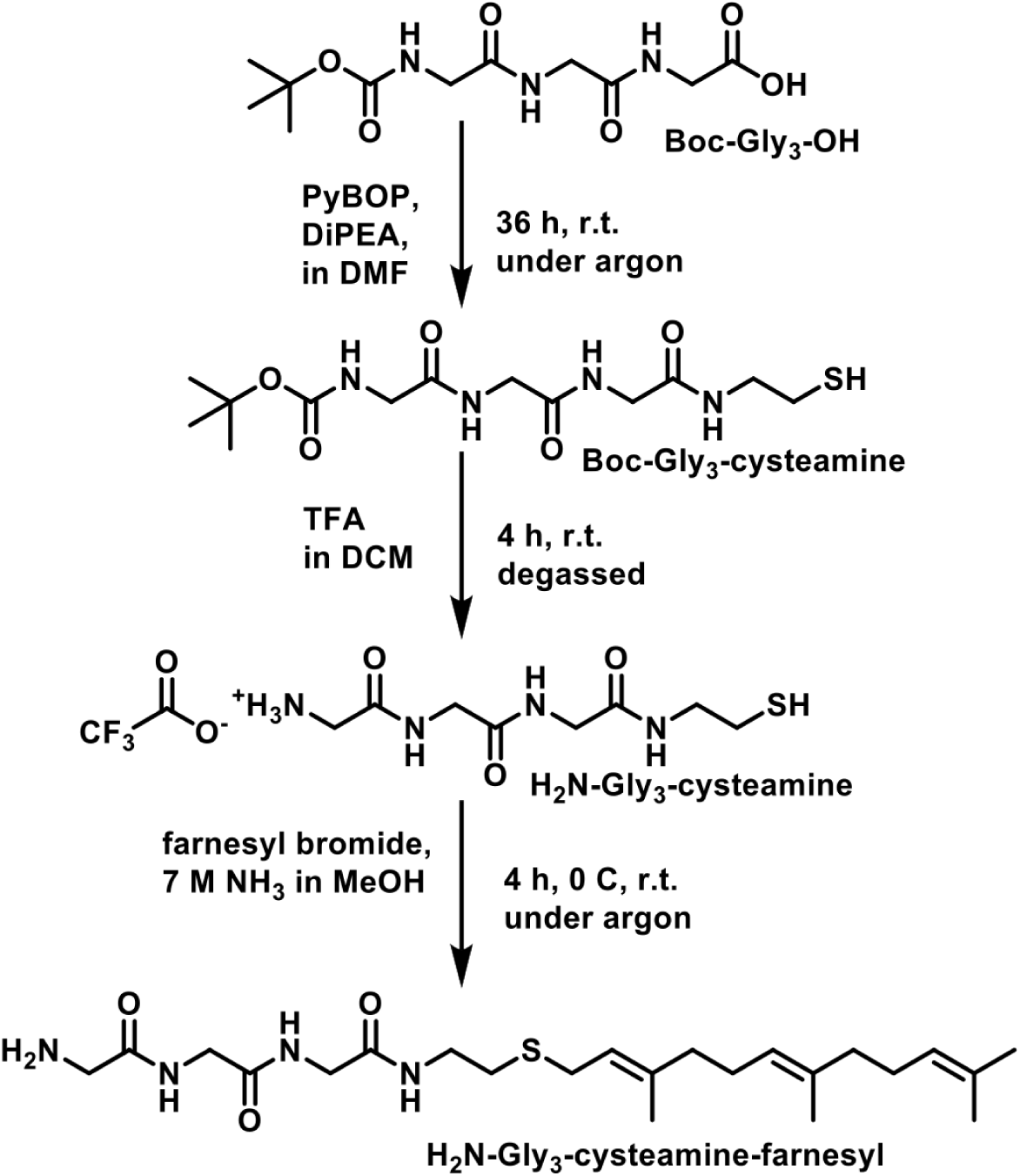
General synthetic procedure going from Boc-Triglycine to the *C*-terminally farnesylated triglycine derivative (’farnesylated triglycine’). Abbreviations: r.t.: room temperature.

The synthesis was carried out according to a modified procedure of Agarwal *et al*. [Agarwal et al., 2015]. Boc-Gly_3_-cysteamine was prepared by dissolving 579.0 mg (2.0 mmol, 1.0 eq.) in 7 mL anhydrous DMF. Then, 1561.0 mg PyBOP (3.0 mmol, 1.5 eq.) and 1293.0 mg (1750 µL, 10.0 mmol, 5.0 eq) DiPEA were added to the clear solution under argon. The solution was stirred for 10 min at room temperature upon which it turned yellow. 455.0 mg (2.0 mmol, 2.0 eq.) cysteamine hydrochloride were added upon which the solution turned colourless again. The reaction mixture was stirred under argon for 36 h at room temperature. The solvent was removed under reduced pressure and the crude, yellow oil was purified via flash column chromatography (DCM/MeOH 9/1) to yield 644.3 mg (93%, 1.85 mmol) of the desired compound as a colourless solid. ^1^H-NMR (400 MHz, DMSO-d6) *δ* = 8.11 (dt, J = 11.9, 5.9 Hz, 2H, NH), 7.92 (q, J = 7.2, 5.9 Hz, 1H, NH), 7.01 (t, J = 6.1 Hz, 1H, NH-Boc), 3.73 (d, J = 5.6 Hz, 2H, CH_2_-Gly), 3.67 (d, J = 5.9 Hz, 2H, CH_2_-Gly), 3.59 (dd, J = 12.0, 6.2 Hz, 2H, CH_2_-Boc), 3.21 (q, J = 6.5 Hz, 2H, CH_2_CH_2_SH), 2.54 (m, 2H, CH2SH, Note: partly overlapping with DMSO signal), 2.39 (t, J = 8.2 Hz, 1H, SH), 1.38 (s, 9H, Boc).

H_2_N-Gly3-cysteamine was prepared by first degassing DCM and TFA separately by sparging with argon for 30 min. 644.0 mg of Boc-Gly_3_-cysteamine (1.85 mmol, 1.0 eq.) were dissolved in 7.4 mL degassed, anhydrous DCM under argon. 1.86 mL degassed TFA were then added at 0^◦^C over the course of 5 min. The solution was then stirred for 4 h at room temperature. The solvent was removed via cold-distillation by subjecting the mixture to a vacuum (approximately 10^−3^ mbar) at 0^◦^C while stirring to yield the NH_2_-Gly_3_-cysteamine TFA salt. The crude was then subjected to a full analysis without further purification. Full conversion was assumed due to the disappearance of the Boc-signal. Note: We found the use of degassed solvents and the removal under reduced pressure at lower temperatures to be essential to avoid oxidation of the free thiol group. Furthermore, it is crucial to remove the Boc-group before introducing the farnesyl residue due to its high lability towards acids [Naider and Becker, 1997]. ^1^H-NMR (400 MHz, DMSO-d6) *δ* = 8.63 (t, J = 5.6 Hz, 1H, NH), 8.26 (t, J = 5.9 Hz, 1H, NH), 8.00 (m, 4H, NH, NH_3_^+^), 3.85 (d, J = 5.7 Hz, 2H, CH_2_-Gly), 3.70 (d, J = 5.7 Hz, 2H, CH_2_-Gly), 3.61 (m, 2H, CH_2_-Gly), 3.26 – 3.16 (m, 2H, CH_2_CH_2_SH), 2.58 – 2.51 (m, 2H, CH_2_SH, Note: partly overlapping with DMSO signal), 2.37 (t, J = 8.2 Hz, 1H, SH).

H_2_N-Gly_3_-cysteamine-farnesyl was synthesised according to a modified procedure of Cini *et al*. [Cini et al., 2009]. 1.85 mmol of H_2_N-Gly_3_-cysteamine were used without further purification. 6.0 mL MeOH were added to dissolve H_2_N-Gly_3_-cysteamine. 7.9 mL of 7M NH_3_ in MeOH were added dropwise at 0^◦^C under nitrogen while stirring. Subsequently, 428.0 mg farnesyl bromide (407 µL, 1.5 mmol, 1.0 eq.) were added. It was stirred for 3 h at 0^◦^C and for 1 h at room temperature. The solvent was removed under reduced pressure at room temperature. The yellow residue was suspended in 15 mL H2O and washed with 1-butanol three times (15 mL + 5 mL + 5 mL). The combined organic phases were dried over MgSO_4_ and filtered under a stream of argon. The solvent was removed under reduced pressure at room temperature. Note: The compound was directly subjected to analysis and no further purification was attempted due to the labile nature of the farnesyl residue which is well established [Naider and Becker, 1997]. The spectra showed some impurities of residual 1-butanol, water and a minor impurity in the aromatic region of unknown origin. ^1^H-NMR (400 MHz, DMSO-d6) *δ* = 8.55 (s, 1H, NH), 8.22 (d, J = 6.4 Hz, 1H, NH), 7.96 (s, 1H, NH), 5.24 – 5.13 (m, 1H, CH=C), 5.06 (d, J = 5.9 Hz, 2H, CH=C), 3.80 (d, J = 3.9 Hz, 2H, CH_2_-Gly), 3.68 (d, J = 5.7 Hz, 2H, CH_2_-Gly), 3.48 (s, 2H, CH_2_-Gly), 3.22 (q, J = 6.9 Hz, 2H, CH_2_CH_2_S), 3.14 (d, J = 7.7 Hz, 2H, CHCH_2_CH_2_S), 2.47 (m, 2H, CH_2_SH, Note: partly overlapping with DMSO signal), 2.10 – 1.97 (m, 6H, CH_2_-farnesyl), 1.91 (dd, J = 15.6, 7.9 Hz, 2H, CH_2_-farnesyl), 1.65 (s, 6H, CH_3_-farnesyl), 1.56 (s, 6H, CH_3_-farnesyl).

The molar mass of the farnesylated triglycine (452 g/mol) was confirmed through mass analysis by Max Perutz Labs Vienna (S7).

### Condition screens for Sortase-mediated reactions

To easily optimize the reaction conditions for Sortase-mediated reactions, Cdc42 was labeled at the *C*-terminus with Alexa488 peptide in a Sortase-mediated reaction [Guimaraes et al., 2013]. To obtain Alexa488 peptide, Alexa Fluor^TM^ 488 C_5_ Maleimide, Invitrogen) was ligated to Gly-Gly-Gly-Gly-Gly-Cys peptide (Biomatik) in a 1:2 molar ratio, as described previously [Liu et al., 2018, Nanda and Lorsch, 2014].

Sortase A (Octamutant, BPS Bioscience) was incubated with Cdc42 and Alexa488-peptide at stated molar ratio in labeling buffer and incubated for stated time points at 4^◦^C or room temperature (see S2). Labeling efficiency was analyzed by SDS-PAGE imaged using a Cy2 filter (Amersham Typhoon Biomolecular Imager, Cytiva).

### Sortase-mediated farnesylation

Cdc42 (Flag-Cdc42-His, His-Cdc42-Flag) was labeled at the *C*-terminus with farnesylated triglycine in a Sortase-mediated reaction [Guimaraes et al., 2013].

#### Labeling of Flag-Cdc42-His

Sortase A (Octamutant, BPS Bioscience) was incubated with Cdc42 and farnesylated triglycine (dissolved in DMSO) at a 2:100:10’000 molar ratio in labeling buffer (204 mM Tris-HCl (pH=7.5), 100 mM NaCl, 10 mM MgCl_2_, 20 mM CaCl_2_, 1mM 2-Mercaptoethanol, 3 mM GTP, 1% Brij58 (Surfact-Amps 58, Thermo Scientific), supplemented with 2 mM freshly prepared dithiothreitol; with a final DMSO content of 22%) for 72 h at 4^◦^C. The sample was spun for 5 min at 13000× g, after which precipitated protein was removed. Reaction mixtures were gel-filtrated on a HiPrep 16/60 Sephacryl S-300 HR column (Cytiva) equilibrated with SEC buffer that was supplemented with 0.1% Brij58 (Surfact-Amps 58, Thermo Scientific). Peak fractions were pooled and immediately loaded onto a HisTrap^TM^ excel column (Cytiva). The flow-though was loaded again and this cycle was repeated for a total of three times. After the last cycle the flow-through, containing the final product, was concentrated (Amicon Ultra Centrifugal Filter, 10 kDa MWCO, Millipore) and flash frozen in liquid nitrogen for storage.

#### Labeling of His-Cdc42-Flag

Sortase A (Octamutant, BPS Bioscience) was incubated with Cdc42 and farnesylated triglycine (dissolved in DMSO) at a 2:100:2’000 molar ratio in labeling buffer (236 mM Tris-HCl (pH=7.5), 100 mM NaCl, 10 mM MgCl_2_, 20 mM CaCl_2_, 1mM 2-Mercaptoethanol, 0.1 mM GDP/GTP, 2% CHAPS (Roche), supplemented with 2 mM freshly prepared dithiothreitol; with a final DMSO content of 6%) for 72 h at 4^◦^C. The sample was spun for 5 min at 13000× g, after which precipitated protein was removed. Reaction mixtures were gel-filtrated on a Superdex 75 Increase 10/300 column (Cytiva) equilibrated with HIC loading buffer. Peak fractions were pooled and immediately loaded onto a HiTrap^TM^ Butyl HP column (Cytiva) equilibrated with HIC loading buffer. After several rounds of washing (using HIC washing buffer), the protein was eluted in a gradient elution (using HIC elution buffer and HIC washing buffer). Peak fractions were flash frozen in liquid nitrogen for storage.

### Mass analysis of farnesylated Cdc42

For mass analysis, samples were dialyzed in (a) SEC buffer, (b) SEC buffer supplemented with 0.1% *n*-Dodecyl *β*-D-maltoside (DDM, Thermo Scientific), and (c) SEC buffer supplemented with 0.1% CHAPS (Roche). Samples were analyzed using intact protein mass-spectrometry analysis, carried out by Max Perutz Labs Vienna: Protein samples were diluted directly to 10 ng/µL with H_2_O or in case of reduction diluted 1:10 with 1 mM TCEP, incubated for 30 min at RT and then diluted to 10 ng/µL. Up to 2 µL (20 ng) were loaded on an XBridge Protein BEH C4 column (2.5 µm particle size, dimensions 2.1 mm × 150 mm; Waters) using a Vanquish^TM^ Horizon UHPLC System (Thermo Fisher Scientific) with a working temperature of 50^◦^C, 0.1% formic acid as solvent A, 99.92% acetonitrile and 0.08% formic acid as solvent B. The gradient started with 12% solvent B, ramped in 5 min to 40% solvent B, in 1 min from 40% to 64% solvent B and in 4 min from 64% to 72% solvent B at a flow rate of 250 µL/min. The eluting proteins were analyzed on a Synapt G2-Si coupled via a ZSpray ESI source (Waters). MS1 spectra were recorded with MassLynx V 4.1 software (Waters) in positive polarity and resolution mode in a m/z range from 600 – 2000. Every 30 s the peptide GluFib was recorded over the LockSpray and used for internal calibration of the spectra. Data were analyzed using the MaxEnt 1 process to reconstruct the uncharged average protein mass.

### Liposome co-floatation assay

Protocol adapted from [Meca et al., 2019]. Lipid stocks of 10 mg/ml in chloroform were mixed into 200ul chloroform in molar ratio of 80 DOPC : 20 DOPS (DOPC: 1,2-dioleoyl-sn-glycero-3-phosphocholine, DOPS: 1,2-dioleoyl-sn-glycero-3-phospho-L-serine, Avanti Polar Lipids) to a final concentration of 5mg/ml in 300µL 0% sucrose buffer (20 mM Tris (pH = 7.5), 150mM NaCl). The lipid solution was then gently dried under a N_2_ air stream and left in a desiccated for 1 hour. Lipids were then rehydrated with 300µL liposome buffer (20mM Tris (pH = 7.5), 150mM NaCl) by vortexing three times for 30 s followed by a 30 resting period. Five cycles of freeze-thaw were applied in liquid nitrogen alternated with a 45^◦^C water bath after which the lipids were bath sonicated for 15 min. After liposome formation, 0.1% Brij58(Surfact-Amps 58, Thermo Scientific) was added to the mixture to match the detergent concentration of the protein samples. GTP is then added to the protein to a final concentration of 300mM GTP, which is about 10,000-fold excess with the protein. All protein was for this assay was handled with Protein LoBind® Tubes (Eppendorf) and Low Binding TripleA pipette tips (Westburg) to prevent protein loss due to the hydrophobic prenylation sticking to the plastics. 17 µL of the protein-GTP mixture was then added to 38µL of liposomes. A small scoop, about one third of the total sample volume, was placed in the tubes and left overnight on a turntable at 4^◦^C to take up the detergent. 40 µL of the mixture was carefully pipetted out, without taking the biobeads, and incubated for 30 minutes at 30^◦^C to promote protein-liposome binding. Then a 60% sucrose buffer (20mM Tris (pH = 7.5), 150mM NaCl, 60% w/v sucrose) was mixed in to a final sucrose content of 30% w/v. This was overlayed by 80µL 25% sucrose buffer (20mM Tris (pH = 7.5), 150mM NaCl, 25% w/v sucrose) and finally overlayed by 80µL 0% sucrose buffer (20mM Tris (pH = 7.5), 150mM NaCl) resulting in a sucrose density gradient with three layers. The samples were then centrifuged for 30 minutes at 42,000 rpm at 4^◦^C. After centrifugation (Optima L-90K, Beckman Coulter), the three layers were separated through carefully pipetting 80µL from the top of the sample per fraction and mixing with SDS loading buffer (Laemmli buffer, [Laemmli, 1970]) for further analysis on SDS-gels and anti-Cdc42 blots. Due to the low concentration of protein in the final sample and protein getting lost due to hydrophobic interactions with the used plastics, the SuperSignal West Femto detection kit(Thermo Scientific) was used for activation diverging from the protocol described below.

### SDS-PAGE

Proteins were analyzed using precast gradient gels (Mini-PROTEAN TGX Stain-Free Protein Gels, Bio-Rad) and freshly prepared SDS-PAGE gels (12-15% acrylamide). In brief, a solution of 375 mM Tris-HCl (pH=8.8), 30-37.5 v/v% 40% acrylamide solution (Biorad), 0.2 w/v% sodium dodecyl sulfate (Sigma Aldrich), 0.5 v/v% 2,2,2-Trichloroethanol (Sigma Aldrich), 0.1 w/v% ammonium persulfate (Sigma Aldrich), and 0.1 v/v% N,N,N’,N’-tetramethyl ethylenediamine (Sigma Aldrich) was prepared and casted into 1.00 mm mini-protean glass plates (Biorad), filling them up to 80%. To protect the gel surface from drying, a layer of isopropanol (Sigma Aldrich) was added. The gel was let solidify for 20 min, after which the isopropanol layer was removed. A solution of 155 mM Tris-HCl (pH=6.5), 10 v/v% 40% acrylamide solution (Biorad), 0.2 w/v% sodium dodecyl sulfate (Sigma Aldrich), 0.1 w/v% ammonium persulfate (Sigma Aldrich), and 0.1 v/v% N,N,N’,N’-tetramethyl ethylenediamine (Sigma Aldrich) was prepared and added to the existing gel layer, after which a well comb (Biorad) was added. The gel was let solidify for 20 min.

Cell and protein samples were mixed with SDS loading buffer (Laemmli buffer, [Laemmli, 1970]). Before loading onto the SDS-PAGE gels, they were kept for 5 min at 95^◦^C. Gels were run for 5 min at 130 V followed by 55min min at 180 V (PowerPac Basic Power Supply (Biorad)). Imaging was done on a ChemiDoc MP (Biorad) using the ‘Stain-free gels’ feature and automatic exposure time determination. Precision Plus Protein Unstained standard (Biorad) was used as a protein standard.

### Western blotting

After SDS-PAGE, the sample was transferred from the SDS-PAGE to a blotting membrane (Trans-Blot Turbo Transfer Pack, Bio-rad) using the ‘Mixed MW’ program of the Trans-Blot Turbo Transfer System (Bio-rad). The blotting membrane was incubated with Immobilon signal enhancer (Millipore) at 4^◦^C for 18 h. The blotting membrane was incubated with primary antibody, diluted in Immobilon signal enhancer (Tab. 3), at room temperature for 1 h. It was washed thrice with TBS-T (10 mM Tris-HCl (pH=7.5), 150 mM NaCl, 0.1 v/v% Tween-20 (Sigma Aldrich)). For each washing step the blotting membrane was incubated with TBS-T at room temperature for 20 min. The blotting membrane was incubated with secondary antibody, diluted in Immobilon signal enhancer (Tab. 3), at room temperature for 1 h, after which it was again washed thrice with TBS-T. SuperSignal West Pico Mouse IgG Detection Kit (Thermo Scientific) was used for activation. Imaging was done on a ChemiDoc MP (Biorad) using the ‘Chemi Sensitive’ feature and automatic exposure time determination. Precision Plus Protein Unstained standard (Biorad) and Precision Plus Protein All Blue Standard (Biorad) were used as protein standards.

**Table 3.**
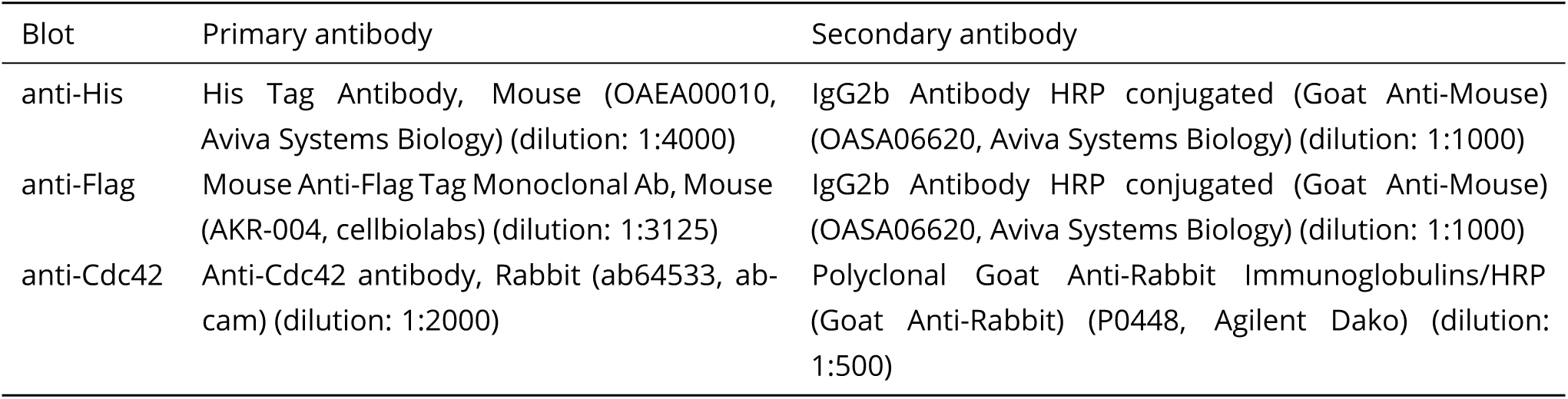
Antibodies used for Western blotting.

| Blot | Primary antibody | Secondary antibody |
| --- | --- | --- |
| anti-His | His Tag Antibody, Mouse (OAEA00010, Aviva Systems Biology) (dilution: 1:4000) | IgG2b Antibody HRP conjugated (Goat Anti-Mouse) (OASA06620, Aviva Systems Biology) (dilution: 1:1000) |
| anti-Flag | Mouse Anti-Flag Tag Monoclonal Ab, Mouse (AKR-004, cellbiolabs) (dilution: 1:3125) | IgG2b Antibody HRP conjugated (Goat Anti-Mouse) (OASA06620, Aviva Systems Biology) (dilution: 1:1000) |
| anti-Cdc42 | Anti-Cdc42 antibody, Rabbit (ab64533, abcam) (dilution: 1:2000) | Polyclonal Goat Anti-Rabbit Immunoglobulins/HRP (Goat Anti-Rabbit) (P0448, Agilent Dako) (dilution: 1:500) |

## Acknowledgements

We thank F. van Opstal and R. van der Valk for experimental assistance, and S. Farooq her pioneering work on Cdc42 membrane binding, and N. Dekker (TU Delft) for the plasmid pET28a-His-mcm10-Sortase-Flag. We thank the group of P. Schwille (MPI Martinsried) for sharing negative results, and the group of A. Jakobi (TU Delft) for assistance in exploring the *E. coli*-based farnesylation method. We thank Régis Lemaitre, Aliona Bogdanova, and Eric Geertsma from the Protein Biochemistry Facility (MPI-CBG in Dresden) for the help in the exploration of prenylation of Cdc42 in insect cells. Intact protein mass analysis was performed by the Mass Spectrometry Facility at Max Perutz Labs using the VBCF instrument pool.

## Contributions

S. Tschirpke: Conceptualization, Methodology, Investigation, Formal analysis, Validation, Writing - Original Draft, Writing - Review & Editing, Visualization, Project administration. N. Hettema: Methodology, Investigation, Formal analysis, Validation, Writing - Original Draft, Writing - Review & Editing, Visualization. B. Spritzbarth: Methodology, Investigation, Formal analysis, Validation, Writing - Original Draft, Visualization. R. Eelkema: Conceptualization, Methodology, Investigation, Funding acquisition, Project administration, Writing - Review & Editing. L. Laan: Conceptualization, Methodology, Investigation, Funding acquisition, Project administration, Writing - Review & Editing.

## Supplement S1

We reviewed the literature for methods to obtain membrane-binding Cdc42 (S1 Fig. 1). Of these, we unsuccessfully attempted:

- Expression of Cdc42 in yeast cells [Das et al., 2012] (S1 Fig. 1a)
- Expression of Cdc42 in insect cells [Zheng et al., 1994, Zheng et al., 1995] (S1 Fig. 1b)
- Co-expression of the FTase and Cdc42 in *E. coli* (established previously for GBP1 [Fres et al., 2010]) (S1 Fig. 1c)
- Cdc42 constructs with alternative membrane binding domains [Meca et al., 2019, Bendezú et al., 2015] (S1 Fig. 1g)

We did not try to utilize *E. coli* cell-free prenylated protein synthesis systems [Sonal et al., 2022] (S1 Fig. 1d) because this method had not yet been published at the time of our study. Further, a significant time investment would have been needed to set up such as system and to establish purification methods for prenylated Cdc42 (which are not described in the publication). While *E. coli* cell-free systems are valuable tools, we believe they are more suited for direct use in microscopy studies rather than as a general method for producing purified prenylated Cdc42.

We also did not pursue *in vitro* geranylgeranylation of bacterial expressed Cdc42 [Golding et al., 2019] (S1 Fig. 1e), as it requires the purification and activity testing of another enzyme (GGTase-I). In retrospect, considering the challenges we faced with other methods, this approach may be worth exploring. In the following, we summarize our progress on methods that ultimately did not (fully) succeed.

**S1 Figure 1.**
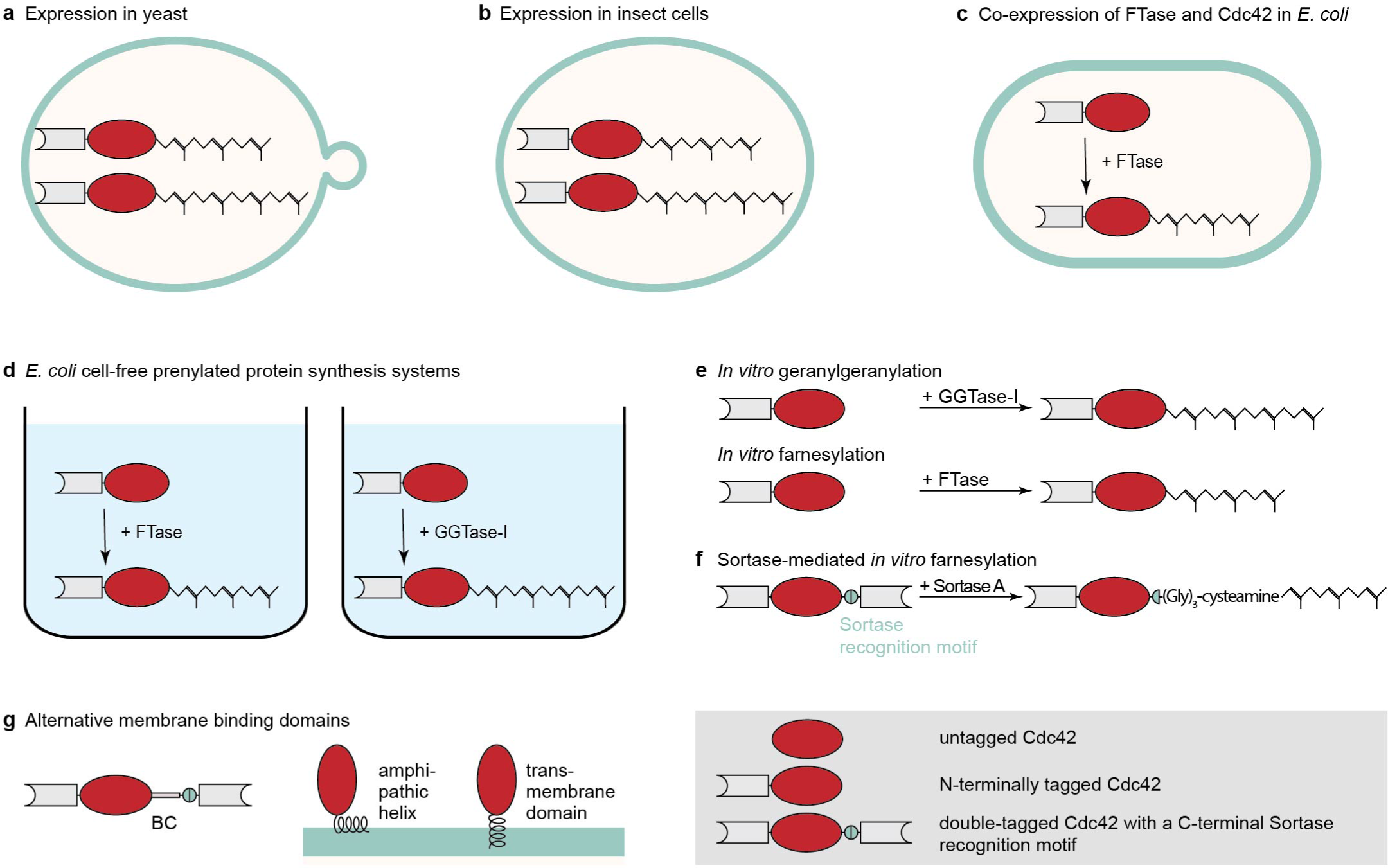
Schematic overview of available methods for obtaining membrane-binding Cdc42. (a) Expression of Cdc42 in yeast cells leads to a mixed pool of farnesylated and geranylgeranylated Cdc42, that can be purified from the membrane fraction [Das et al., 2012]. (b) Expression of Cdc42 in insect cells also results in both prenylation products [Zheng et al., 1994, Zheng et al., 1995]. (c) Co-expression of the FTases and Cdc42 produces farnesylated Cdc42 in *E. coli* (this work, and established previously for GBP1 [Fres et al., 2010]). (d) Use of *E. coli* cell-free prenylated protein synthesis systems [Sonal et al., 2022]. (e) *In vitro* geranylgeranylation or farnesylation of bacterial expressed Cdc42 [Golding et al., 2019]. (f) Sortase-mediated *in vitro* farnesylation of bacterial expressed Cdc42 (this work). (g) Cdc42 constructs with alternative membrane binding domains, such as the basic clusters from Bem1 (BC) (this work, and [Meca et al., 2019]), an amphipathic helix or a transmembrane domain [Bendezú et al., 2015].

### Expression of Cdc42 in yeast cells

We attempted the purification of native Cdc42 from *Saccharomyces cerevisiae*. Cells were lysed using a French press or Freezer mill using lysis buffer (Tab. 2), supplemented with no guanine nucleotide, with 10 mM GDP, or with 10 mM GTP. In all cases the vast majority of protein remained in the pellet fraction, of which it could also not be solubilized using lysis buffer supplemented with 10 mM GTP.

### Expression of Cdc42 in insect cells

To test for Cdc42 prenylation in insect cells, we collaborated with the Protein Biochemistry Facility of MPI-CBG in Dresden. Régis Lemaitre, Aliona Bogdanova, and Eric Geertsma who carried out the following expression and purification tests:

#### Cdc42 constructs

We used two Cdc42 constructs (24 kDa), each containing a N-terminal 10His-tag and 3C cleavage site. One of these was based on yeast cDNA, the other used was codon-optimized for insect cell expression. Additionally, a Cdc42 construct with a sfGFP sandwich-fusion (Cdc42-sfGFP*^SW^*, 52 kDa) [Bendezú et al., 2015, Tschirpke et al., 2023] was created based on yeast cDNA.Their sequences are shown below.

**D7090: HIS10-3C-Cdc42 (using yeast Cdc42 cDNA)**

⇒ Amino acid sequence:

MGSSHHHHHHHHHHSSGRLEVLFQGPAAAMQTLKCVVVGDGAVGKTCLLISYTTNQFPADYVPTVFDNYAVTVMIGDEPYTLGLFDTAGQ EDYDRLRPLSYPSTDVFLVCFSVISPPSFENVKEKWFPEVHHHCPGVPCLVVGTQIDLRDDKVIIEKLQRQRLRPITSEQGSRLARELKA VKYVECSALTQRGLKNVFDEAIVAALEPPVIKKSKKCAIL

⇒ cDNA:

ATGCAAACGCTAAAGTGTGTTGTTGTCGGTGATGGTGCTGTTGGGAAAACGTGCCTTCTAATCTCCTATACAACGAATCAATTTCCAGCC GACTATGTTCCAACAGTGTTCGATAACTATGCGGTGACTGTGATGATTGGTGATGAACCATATACGTTAGGTTTGTTTGATACGGCCGGT CAAGAAGATTACGATCGATTGAGACCCTTGTCATATCCTTCTACTGATGTATTTTTGGTTTGTTTCAGTGTTATTTCCCCACCCTCTTTT GAAAACGTTAAAGAAAAATGGTTCCCTGAAGTACATCACCATTGTCCAGGTGTACCATGCCTGGTCGTCGGTACGCAGATTGATCTAAGG GATGACAAGGTAATCATCGAGAAGTTGCAAAGACAAAGATTACGTCCGATTACATCAGAACAAGGTTCCAGGTTAGCAAGAGAACTGAAA GCAGTAAAATATGTCGAGTGTTCGGCACTAACACAACGCGGTTTGAAGAATGTATTCGATGAAGCTATCGTGGCCGCCTTGGAGCCTCCT GTTATCAAGAAAAGTAAAAAATGTGCAATTTTGTAG

**D7091: HIS10-3C-Cdc42 (using cDNA codon optimized for insect cells)**

⇒ Amino acid sequence: see D7090

⇒ cDNA:

ATGCAAACGTTGAAGTGCGTGGTAGTGGGAGACGGGGCCGTCGGGAAGACCTGTTTACTAATATCTTACACTACGAATCAGTTTCCGGCC GACTATGTCCCCACAGTATTTGATAACTATGCTGTAACCGTTATGATTGGTGACGAACCGTACACCCTGGGCCTCTTCGACACAGCTGGCCAAGAAGATTATGATAGGCTTCGCCCACTCTCCTACCCATCGACTGACGTCTTCTTAGTTTGTTTCTCCGTCATTTCACCACCTAGTTTC GAAAATGTGAAGGAAAAATGGTTTCCGGAGGTACATCACCACTGCCCTGGAGTGCCCTGCTTGGTAGTCGGTACTCAAATCGATCTGCGT GACGATAAAGTGATTATTGAGAAACTTCAGAGACAAAGGCTACGGCCTATCACAAGCGAGCAGGGTAGCCGCCTAGCTAGAGAACTGAAG GCAGTTAAGTATGTAGAGTGCTCTGCCCTCACGCAGCGAGGGCTTAAGAACGTGTTTGATGAAGCAATAGTTGCGGCATTAGAGCCACCC GTTATAAAAAAAAGTAAAAAATGTGCGATCTTGTGA

**D7099: HIS10-3C-cdc42-sfGFP-cdc42 (Cdc42-sfGFP*^SW^*)**

⇒ Amino acid sequence:

MGSSHHHHHHHHHHSSGRLEVLFQGPAAAMQTLKCVVVGDGAVGKTCLLISYTTNQFPADYVPTVFDNYAVTVMIGDEPYTLGLFDTAGQ EDYDRLRPLSYPSTDVFLVCFSVISPPSFENVKEKWFPEVHHHCPGVPCLVVGTQIDLRDDKVIIEKLQRQRLSGGSAMSKGEELFTGVV PILVELDGDVNGHKFSVRGEGEGDATNGKLTLKFICTTGKLPVPWPTLVTTLTYGVQCFSRYPDHMKRHDFFKSAMPEGYVQERTISFKD DGTYKTRAEVKFEGDTLVNRIELKGIDFKEDGNILGHKLEYNFNSHNVYITADKQKNGIKANFKIRHNVEDGSVQLADHYQQNTPIGDGP VLLPDNHYLSTQSVLSKDPNEKRDHMVLLEFVTAAGITHGMDELYKSGPPGRPITSEQGSRLARELKAVKYVECSALTQRGLKNVFDEAI VAALEPPVIKKSKKCAIL

#### Expression tests

All Cdc42 constructs were cloned into a pOCC33 vector and sequence verified. Subsequently, production of baculovirus was performed using the FlexiBac system (as reported in [Lemaitre et al., 2019]), resulting in viruses V7307 (D7090: HIS10-3C-Cdc42 (cDNA)), V7308 (D7091: HIS10-3C-Cdc42 (codon optimized)), and V7309 (D7099: HIS10-3C-cdc42-sfGFP-cdc42 (Cdc42-sfGFP*^SW^*)).

Expression tests were carried out using T. ni (*Trichoplusia ni*) and SF9 (*Spodoptera frugiperda*) cell lines cultivated in suspension in glass culture flasks in ESF 921 serum-free medium (Expression systems). Insect cell cultures, at a density of 1 × 10^6^ cells/ml, were infected with 1% (v/v) of a P2 baculovirus stock and cultivated at 22^◦^C and 28^◦^C. Samples(2mL) were taken at 48 h and 72 h and cells were collected by centrifugation (5 min @ 500 g). Cell pellets were resuspended in 200 µL lysis buffer(2× PBS (pH = 7.5), 2 mM MgCl_2_, 1% (v/v) Triton X100, 1/10’000 (v/v) benzoase, protease inhibitor(Protease Inhibitor Cocktail (EDTA-Free, 100X in DMSO))) and lysed by incubation on ice for 30 min. For SDS-PAGE analysis, a fraction of the crude solution as taken. The remainder was spun down in a bench-top centrifuge (25’000 × g, 30 min, 4^◦^C) and a sample of the supernatant was taken.

Samples of the complete crude cell extract and the supernatant after centrifugation were compared side-by-side using SDS-PAGE and detection by Coomassie-staining. SDS-PAGE analysis suggests that both Cdc42 and Cdc42-sfGFP*^SW^* were expressed, based on the appearance of an additional protein band at the expected molecular weight. Cdc42 expresses only in T. ni, but not in SF9cells. As the Cdc42 band appears only in the crude samples and not in the supernatant, the protein is most likely expressed as insoluble aggregates. No difference in expression level between the Cdc42 construct with yeast cDNA and the construct that was codon optimized for expression in insect cells could be observed (S1 Fig. 2a,b). Expression of Cdc42-sfGFP*^SW^* resulted in green cells. Most color, suggesting the presence of fluorescently labelled protein, went to the pellet after lysis and high speed centrifugation, indicating poor solubility. Fluorescent imaging of SDS-PAGE shows that Cdc42-sfGFP*^SW^*, however, is partially soluble, both in T. ni and SF9 expression conditions. Most signal is observed for T. ni cells for 22^◦^C 48 h and for SF9 cells for 28^◦^C 72 h (S1 Fig. 2c).

#### Purification

T. ni cells were infected with virus V7307 (Cdc42) or V7309 (Cdc42-sfGFP*^SW^*) and grown for 72 h at 22^◦^C, after which the cells were collected by centrifugation. Cells were resuspended in lysis buffer (2× PBS (pH = 7.5), 20 mM imidazole, 1 mM MgCl_2_, 5% glycerol, 1 mM DTT, 1% Trition X100, 1/10’000 benzoase, protease inhibitor), and lysed using a douncer (20 rounds, followed by a 30 min incubation on ice, followed by another 20 rounds). The lysate was clarified by centrifugation (1 h, 23’000 rpm), filtered (0.22 µm), and loaded onto a 1 mL HisTrap column. The column was washed with 20 CV of high salt buffer (6× PBS (pH = 7.5), 50 mM imidazole, 5% glycerol, 1 mM DTT, 0.05% DDM), followed by 10 CV of low salt buffer (2× PBS (pH = 7.5), 20 mM imidazole, 5% glycerol, 1 mM DTT, 0.05% DDM). The protein was eluted with elution buffer (2× PBS (pH = 7.5), 500 mM imidazole, 5% glycerol, 1 mM DTT, 0.05% DDM). Peak fractions were pooled and dialyzed against SEC buffer (2× PBS (pH = 7.5), 5% glycerol, 1 mM DTT, 0.05% DDM), and concentrated. Samples were analyzed by SDS-PAGE (S1 Fig. 3). Concentrated peak fractions were injected onto a SEC column (Superdex 200 increase 10/300), equilibrated in SEC buffer, and eluted using SEC buffer. The elution profile is shown in is shown in S1 Fig. 4 and SDS-PAGE analysis is given in S1 Fig. 5.

Most Cdc42 remained in the pellet after solubilization. Cdc42 represents the main band in the eluate from His-AC,, but is 60% pure based on SDS-PAGE (S1 Fig. 3). In SEC, Cdc42 elutes in one peak close to the void, indicative of aggregated protein (S1 Fig. 4a, S1 Fig. 5).

Cdc42-sfGFP*^SW^* shows a similar behavior: most protein remained in the pellet (observable through the green color), although some remained soluble and could be eluted from His-AC. The protein was of similar purity as Cdc42 (S1 Fig. 3). In SEC, Cdc42-sfGFP*^SW^* elutes in several peaks (S1 Fig. 4b), but most protein comes in broad peaks at very high molecular weights, indicating aggregation (S1 Fig. 5).

These results suggest that the folding state of Cdc42 is compromised; an essential yeast chaperone/interaction partner may be missing. In general, the fact that the protein is misbehaving to such an extend was surprising. So-far, a double band indicative of partial prenylation was not observed. This might also result from the use of gradient gels. However, given the concerns on the folding state of Cdc42, this route was terminated.

**S1 Figure 2.**
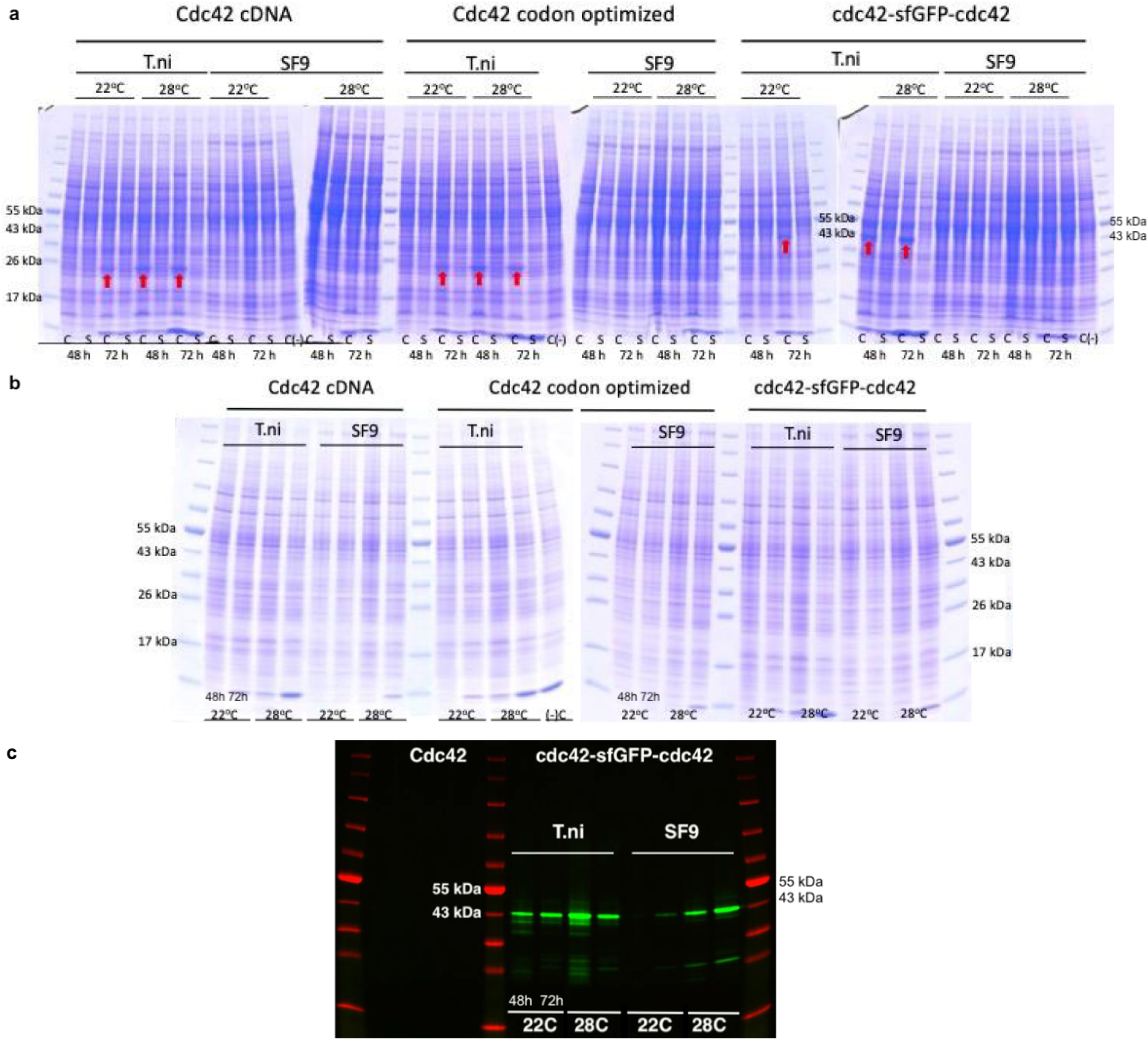
Cdc42 and Cdc42-sfGFP*^SW^* and be expressed T. ni and Sf9 cells, but is hardly soluble. (a) Coomassie-stained SDS-PAGE of the crude cell extract(labeled with ‘C’) and supernatant (labeled with ‘S’) expression test samples. C(-) indicates a negative control containing cells not transfected with any virus. A red arrow indicates the presence of a Cdc42/ Cdc42-sfGFP*^SW^* band. (b) Coomassie-stained SDS-PAGE of supernatant expression test samples. For each condition, the 48 h and 72 h samples are shown. (c) Alexa488-imaged SDS-PAGE of supernatant expression test samples. For each condition, the 48 h and 72 h samples are shown. Cdc42-sfGFP*^SW^* is shown in green and the protein marker is shown in red.

**S1 Figure 3.**
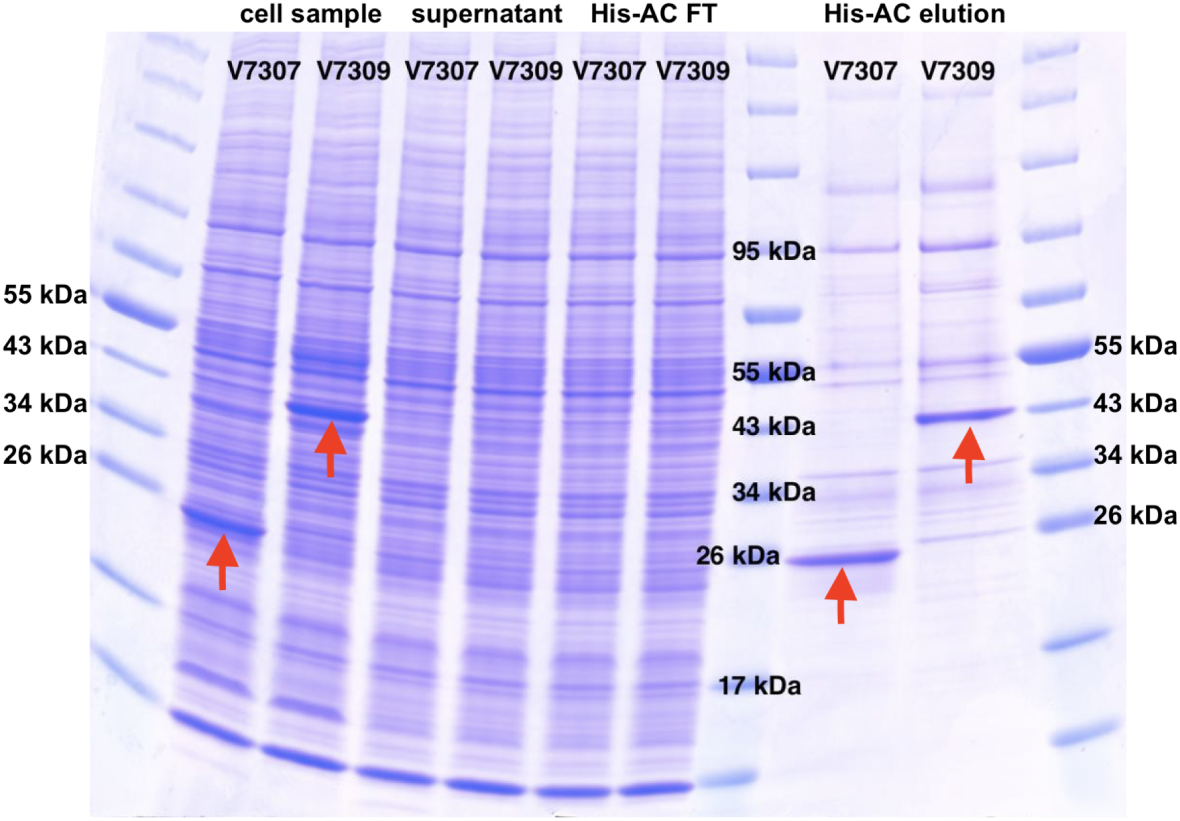
Cdc42 and Cdc42-sfGFP*^SW^* be purified using His-affinity chromatography (His-AC). Coomassie-stained SDS-PAGE of the cell sample, supernatant, His-AC flow-through (FT) and elution of Cdc42 (V7307) and Cdc42-sfGFP*^SW^*(V7309) expressed in T. ni. A red arrow indicates the presence of a Cdc42/ Cdc42-sfGFP*^SW^* band.

**S1 Figure 4.**
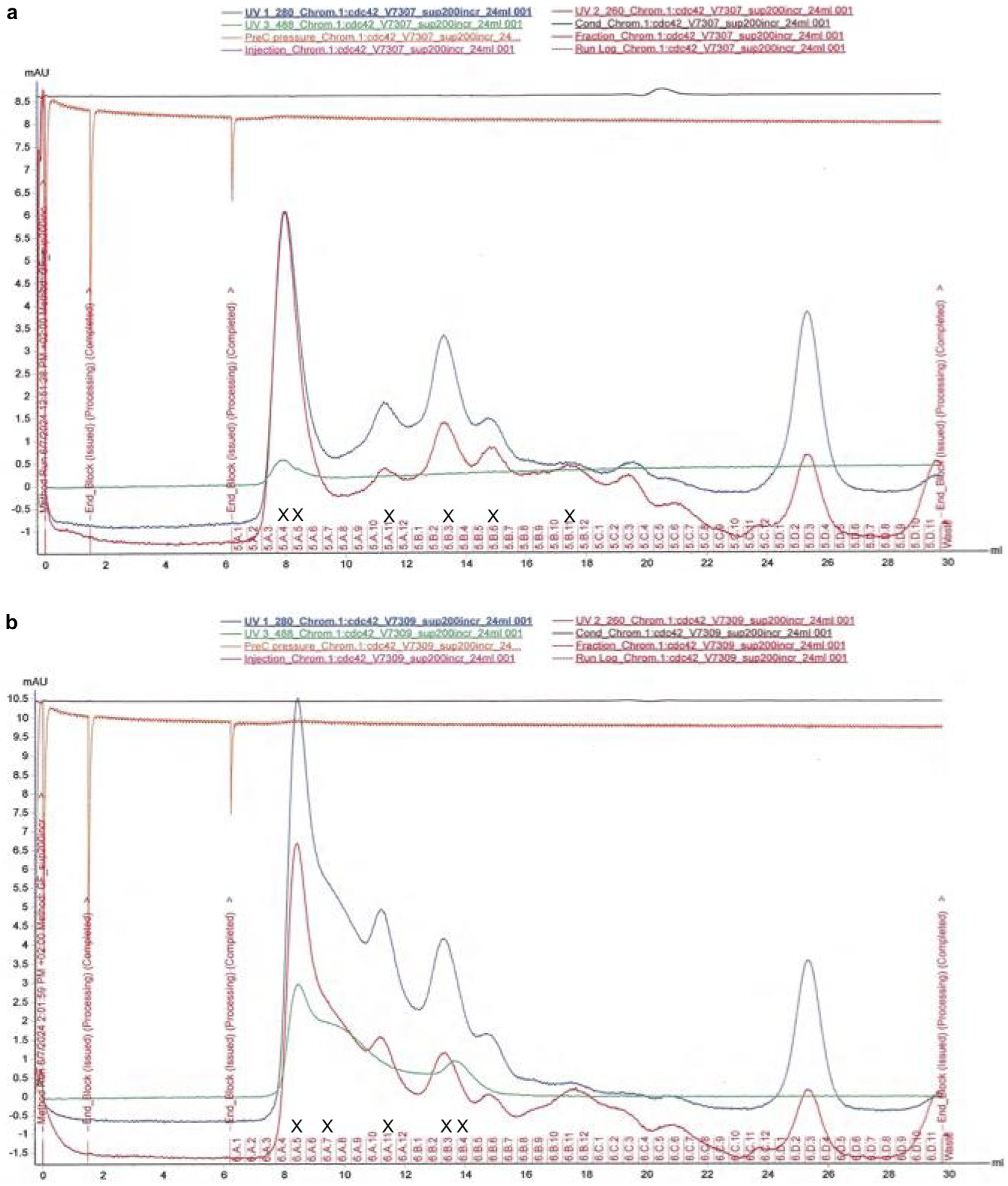
Cdc42 and Cdc42-sfGFP*^SW^* elute close to the void volume in size-exclusion chromatography (SEC). (a) SEC elution profile of Cdc42 (V7307). (b) SEC elution profile of Cdc42-sfGFP*^SW^*(V7309). An ‘X’ indicates the fractions used for SDS-PAGE analysis.

**S1 Figure 5.**
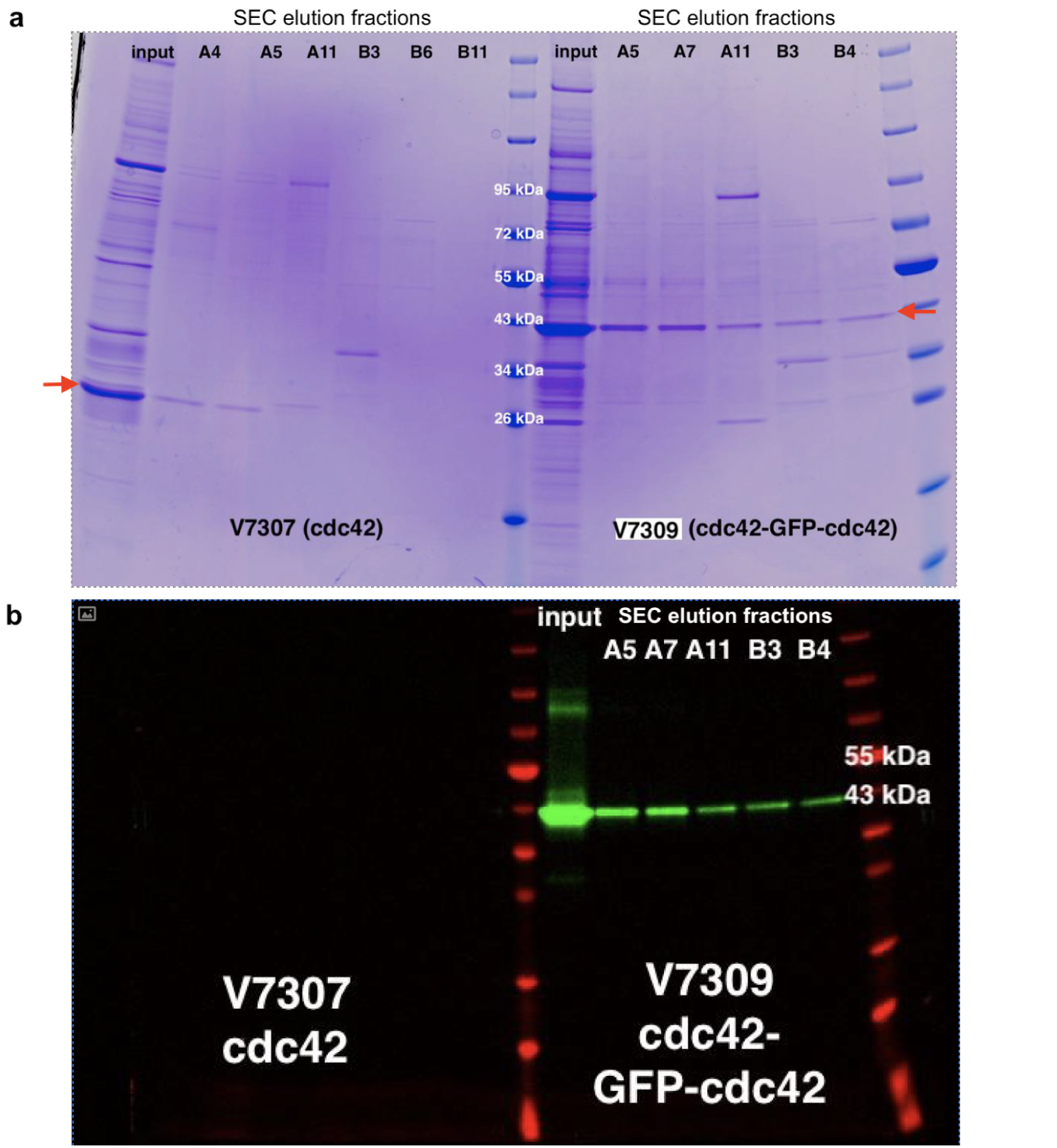
SDS-PAGE analysis of SEC fractions of Cdc42 (V7307) and Cdc42-sfGFP*^SW^*(V7309). (a) Coomassie-stained SDS-PAGE of SEC elution fractions of Cdc42 (V7307) and Cdc42-sfGFP*^SW^*(V7309). A red arrow indicates the size of the Cdc42/ Cdc42-sfGFP*^SW^* band. (b) Alexa488-imaged SDS-PAGE of SEC elution fractions. Cdc42-sfGFP*^SW^* is shown in green and the protein marker is shown in red.

### Co-expression of the FTase and Cdc42 in ***E. coli***

Most proteins, no matter which organisms they originate from, can be expressed and purified from *E. coli*. This bacteria species is easy to manipulate, low in maintenance, grows fast, and lyses under relatively mild conditions. Proteins expressed in a foreign host are also less likely to bind to, and co-purify, one of their natural binding partners. Despite these advantages, the use of bacterial expression systems is in general limited to post-translationally unmodified proteins, as the *E. coli* machinery lacks the enzymes responsible for these modifications. However, attempts have been made to introduce such enzymes into the bacterium [Sugase et al., 2008]. Farnesylation of the human Guanylate-binding protein GBP1 was achieved through co-expression of the *α* and *β* subunit of the human FTase (FTase-*α*, FTase-*β*) in *E. coli* [Fres et al., 2010].

In yeast, farnesylation of Cdc42 is also carried out by an FTase [Gomez et al., 1993]. The substrate specificity of FTases is mainly determined by the sequence of the targets C-terminal CAAX box, and the CAAX sequence preference of mammalian FTases has been found to be nearly identical to that of the yeast enzyme [Caplin et al., 1994, Reid et al., 2004]. Following Fres *et al*., we here also introduced FTase-*α* and FTase-*β* into *E. coli* [Fres et al., 2010] to test if prenylated Cdc42 can be produced by a bacterial expression system (Fig. 1c). We decided to try out farnesylation first, as gernaylgeranylation requires a larger machinery.

Cdc42’s natural CA_1_A_2_X sequence in yeast is CAIL (Fig. 1). The leucine at position four gives the protein a higher affinity for binding to the GGTase-I compared to the FTase [Caplin et al., 1994]. In general, FTases prefer valine (V), isoleucine (I), or leucine (L) on the A_2_ position, and methionine (M), serine (S), or glutamine (Q) on the last position of the CAAX box [Caplin et al., 1994, Hougland et al., 2010, Reid et al., 2004]. In order to optimize for high farnesylation, we designed four Cdc42 constructs with CAAX sequences that matched these criteria and also showed high kinetic values in *in vitro* farnesylation screens with peptide substrates [Caplin et al., 1994, Hougland et al., 2009, Hougland et al., 2010]; CTIS, CAIM, CALQ, CSIM.

#### Expression

To optimize the expression of all involved proteins, we carried out expression screens for (1) the FTase alone, (2) Cdc42 alone, and (3) Cdc42 in presence of the FTases, and analysed expression levels through anti-His Western blotting ^3^ (S1 Fig. 6). Three expression conditions were chosen: ‘f’: a strong and fast expression at elevated temperatures, induced by a high amount of IPTG (3 h at 37^◦^C with 1 mM IPTG). ‘s’: a low and slow expression at lower temperatures, induced by a smaller amount of IPTG (18 h at 18^◦^C with 0.2 mM IPTG). ‘AI’: a self-inducing combined approach (3 h at 37^◦^C + 18 h at 18^◦^C) [Studier, 2005].

We tested the expression levels of the FTase by 6His-tagging the FTase-*α* subunit (His-FTase-*α*) and co-expressing it with untagged FTase-*β* (S1 Fig. 6a). All conditions show a band at ∼50 kDa in the anti-His Western blot, corresponding to His-FTase-*α* (45 kDa). One or two lower bands are visible as well, likely representing products of degradation or of prematurely terminated translations of FTase-*α*. The expression of FTasecan not be assessed (as FTase-*β* is not 6His-tagged), but we assume they are equal to that of FTase-*α*. These results indicate that the FTase expresses well in all conditions.

We tested how N-terminally 6His-tagged Cdc42 constructs with farnesylation-optimized CAAX sequences express. As a control, we utilized His-Cdc42:CAIA, a Cdc42 construct with the CAAX sequence CAIA that does not favor farnesylation. In absence of FTase, all Cdc42 constructs (26 kDa) express well - intense bands at 25 kDa are visible in the anti-His Western blot^4^ in all conditions (S1 Fig. 6c). The expression levels seem mostly unaffected by the specific CAAX sequences.

Next, we investigated the effects of co-expression of Cdc42 and FTase. Less Cdc42-CTIS expresses in presence of FTase compared to absence of FTase (S1 Fig. 6b). This is both true when the 6His-tagged and the untagged FTase versions are used. Of the tested expression conditions, ‘f’ seem to produce the most Cdc42 (both in absence and presence of FTase). The blots of His-Cdc42:CTIS in presence of His-FTase-*α*/FTase-*β* show again two bands at 50 and 40 kDa, corresponding to His-FTase-*α*. Thus, co-expression of Cdc42:CTIS and FTase in *E. coli* is viable. We examined the effect of Cdc42’s CAAX sequence on protein expression levels in cells with both Cdc42 and His-FTase-*α*/FTase-*β* (S1 Fig. 6d) or FTase-*α*/FTase-*β* (S1 Fig. 6e). The blots show again bands for Cdc42 and for His-FTase-*α* (S1 Fig. 6d), illustrating that co-expression of His-FTase-*α*/FTase-*β* and all Cdc42 constructs is possible. The intensity of the His-FTase-*α* bands is not influenced by the CAAX sequence of Cdc42. Hence, Cdc42 does not influence the expression of FTase. We assume that this also applies for the untagged FTase and Cdc42. However, the distinct CAAX sequences seem to influence Cdc42 expression levels: In presence of tagged and untagged FTase-*α*/FTase-*β*, His-Cdc42:CAIA expresses best in condition ‘f’, and His-Cdc42:CSIM and His-Cdc42:CTIS express best in condition ‘s’ and ‘AI’ (S1 Fig. 6d)^5^.

As farnesylation levels are not known, it is difficult to predict if the differences in Cdc42 expression in presence of FTase correspond to differences in Cdc42 farnesylation levels. It is surprising that the CAAX sequences are only influencing Cdc42 expression levels in presence of FTases and that they do not influence the expression of FTase. FTase has a low affinity for the CAAX sequence CAIA. We thus expect His-Cdc42:CAIA to not, or only to a very limited extend, get farnesylated. In presence of FTase, a very small (S1 Fig. 6e, condition ‘AI’) or a medium amount (S1 Fig. 6d, condition ‘AI’) of His-Cdc42:CAIA expresses and more of His-Cdc42:CTIS expresses. If there is a correlation between expression levels and farnesylation, we therefore would assume that high expression levels mean high farnesylation (otherwise we would expect His-Cdc42:CAIA to express way stronger than all other Cdc42 constructs).

#### Farnesylation

We expressed His-Cdc42:CSIM (condition ‘s’ and ‘AI’) and His-Cdc42-CTIS (condition ‘AI’), as they exhibited a high expression level in ‘AI’ in presence of FTase (S1 Fig. 6e), and purified Cdc42 using His-AC. To perform mass-spectrometry, the protein was dialyzed, precipitated with chloroform methanol, and digested with GluC, as described previously [Wessel and Flügge, 1984, Fres et al., 2010].

Of all purified His-Cdc42:CSIM (condition ‘s’ and ‘AI’), our initial LC-MS analysis suggested that less than 0.1% was farnesylated. The sample of His-Cdc42:CTIS (condition ‘AI’) precipitated during dialysis. In the soluble fraction less than 0.1%, and in protein in the pellet 5% was farnesylated, yielding in total less than 1% farnesylated protein. These results suggest that the CAAX sequence indeed influences farnesylation, but that the Cdc42 expression levels might not that strongly correlate with farnesylation levels. However, follow-up LC-MS analysis revealed that the obtained results are not reproducible/ not reliable, likely due to peptide loss during processing and uneven digestion, as GluC appeared to have preferred cleavage sites. As a result, we are uncertain about the extent of Cdc42 farnesylation.

#### Outlook

Our data shows that in principle *E. coli* can be engineered to produce farnesylated Cdc42, but it remains unclear how effective this approach is.

If the yield is low, as our data suggests, further optimization will be be necessary. So-far we only purified and analysed the soluble protein fraction. If farnesylation is occurring, most farnesylated Cdc42 might as well bind to membranes and be in the membrane fraction after lysis. It may be worthwhile to investigate whether Cdc42 is localized in the membrane fraction and how much of it is farnesylated.

What other optimizations are possible? We already altered Cdc42’s CAAX sequence to make it more farnesylation prone. Directly upstream of the CAAX box is the polybasic region (PBR), which consists of five mostly basic amino acids (aa). The PBR is a common feature of prenylated GTPases and affects the FTase’s affinity for the protein [Hicks et al., 2005, Williams, 2003]. Even though the PBR is a common feature, its sequence varies among GTPases. For example, *S. cerevisiae* Cdc42 has the PBR sequence KKSKK (Fig. 1) and that of human Cdc42 is KKSRR. Here we integrated human FTase in *E. coli* to farnesylate yeast Cdc42. The CAAX sequence preferences of yeast FTase have been found to be nearly identical to the preferences of the mammalian enzyme [Caplin et al., 1994, Reid et al., 2004], suggesting that the protein origin is not that important. The human and yeast PBR are also quite similar. However, single mutations in the PBR have been shown to have the capacity to significantly influence protein behavior [Zhang et al., 1999]. To test if this is the case for FTase’s affinity for yeast Cdc42, a Cdc42 version with the PBR of human Cdc42 (KKSRR) could be generated.

Another reason for a low farnesylation yield could be that the CAAX box is not that accessible to the FTase. It might be worthwhile to generate Cdc42 versions where a short linker is placed either upstream of the PBR or in-between PBR and CAAX box, or where the C-terminal region of Cdc42 (PBR and CAAX box) is replaced by the C-terminal region of the GTPase GBP1. The latter approach alters Cdc42 the most, but GBP1 has been successfully and reproducibly been farnesylated in *E. coli* [Fres et al., 2010, Kuhm et al., 2023]. It thus might give most insight into whether the C-terminal region of Cdc42 is hindering farnesylation and if farnesylation of Cdc42 in *E. coli* is effective.

#### LC-MS analysis

LC-MS analysis was performed using a Agilent LC/MS system consisting of a high pressure liquid chromatography set-up coupled to a triple-quadrupole (QQQ) mass spectrometer (G6460C) equipped with a standard electrospray ionization (ESI) source. Both systems were operated through MassHunter data acquisition software (version 10.1). The digested peptides were acidified with 25mM formic acid and delivered to a CSH C18 guard-column and a CSH C18 column (Waters) (2.1 mm by 50 mm, 1.7-µm pore size) at 40^◦^C with a flow rate of 0.5 ml/min using the following binary gradient: 5% buffer B (acetonitrile, 25 mM formic acid), ramp to 95% B in 8 min followed by a 2 min hold at 95% buffer B (acetonitrile, 25 mM formic acid), 2 min ramp back to 5% buffer B (acetonitrile, 25 mM formic acid) and 3 min re-equilibration buffer A, (Milli Q, 25mM formic acid). Next, peptides were eluted from the column and the eluent was sprayed into the mass spectrometer operated in data-dependent mode, as in dynamic multiple-reaction monitoring (dMRM) mode using transitions. The transitions were generated from protein sequences with Skyline [MacLean et al., 2010]. The modified peptides were selected from *in silico* digested peptides based on prior knowledge of the modification location in their sequences. Non-modified peptides were selected based on their length (7-25 amino acids). Each MRM transition was optimized for the fragmentor voltage and collision energy. The dMRM was acquired in positive mode with a cycle time of 500 ms.

**S1 Figure 6.**
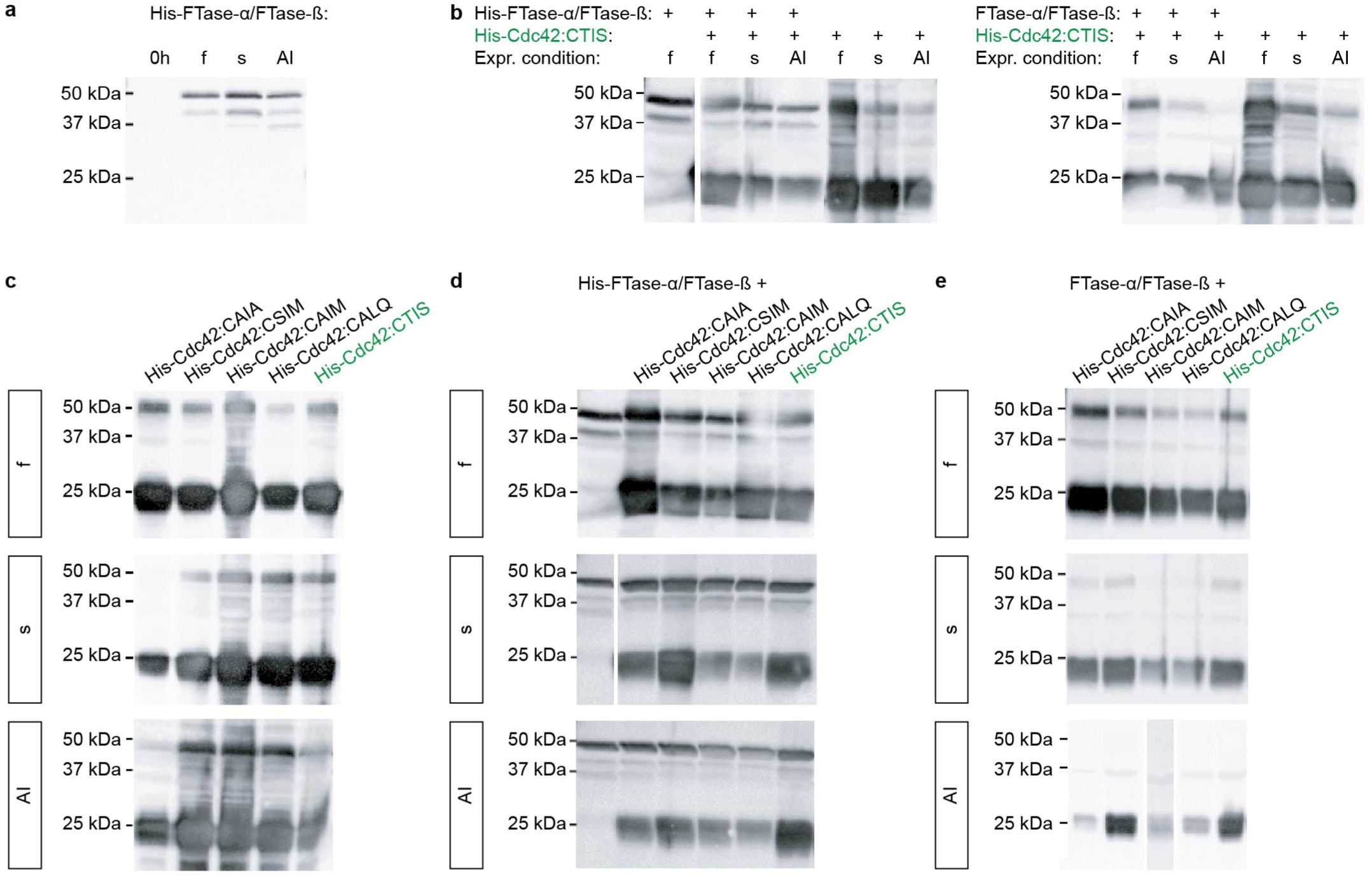
Farnesylation of Cdc42 in ***E. coli***: both Cdc42 and FTases express robustly. Expression screens (condition ‘f’, ‘s’, and ‘AI’), monitored by anti-His Western blotting, of (a) His-FTase-*α* in presence of untagged FTase-*β*, (b) His-Cdc42:CTIS in presence and absence of His-FTase-*α*/FTase-*β* (left) or FTase-*α*/FTase-*β* (right), (c) Cdc42 constructs with different CAAX sequences, (d) Cdc42 constructs with different CAAX sequences in presence of His-FTase-*α*/FTase-*β*, and (e) Cdc42 constructs with different CAAX sequences in presence of FTase-*α*/FTase-*β*.

### Use of Cdc42 constructs with alternative membrane binding domains

The farnesyl or geranylgeranyl tail in Cdc42 is mainly responsible for anchoring the protein to the membrane [Peurois et al., 2018]. If membrane-binding is the main objective of adding a post-translational modification to the protein, any other membrane-binding modification may also be sufficient. Earlier work in fission yeast by Bendezu *et al*. showed that cells in which prenylated Cdc42 got replaced with a Cdc42 allele with a prenylation independent membrane binding mechanism (Cdc42 with a transmembrane domain from the protein Psy1, or Cdc42 with an amphipathic helix from the protein Rit) polarized and showed viability [Bendezú et al., 2015]. We considered testing if such constructs are viable alternatives for prenylated Cdc42 for in vitro experiments, but disregarded them, as proteins with transmembrane domains are typically difficult to purify. Prior attempts to overexpress Cdc42 with the RitC C-terminal amphipathic helix [Bendezú et al., 2015] in bacterial cells yielded misfolded protein pellets which could not be solubilized by standard urea-based refolding protocols (data by P. Schwille group, MPI Martinsried).

#### Cdc42-BC

Recently, Meca *et al*. introduced another membrane-binding Cdc42 construct: here, the membrane-binding property originates from the so-called basic cluster (BC) region of the yeast protein Bem1 [Meca et al., 2019]. The BCs are a 23 to 74 amino acid (aa) long unstructured region of mostly positively charged aa, that are responsible for anchoring Bem1 to negatively charged membranes *in vitro*. It was shown that cells containing a fusion of Cdc42 with the first part of the BC region (26 aa) are viable and polarize, suggesting that the membrane binding capability of the BC region is sufficient to mimic those of the prenyl group. Adding the BCs as membrane binding domain, rather than adding a transmembrane domain, might also be advantageous in other ways: (1) Once Cdc42 is fully folded, its N-terminus and C-terminus are in close proximity to each other. N-terminal fusions of Cdc42 and fluorescent proteins lead to not fully functional proteins [Bendezú et al., 2015]. It is possible that the C-terminal addition of any structural domain, such as an amphipathic helix, might have a similar effect on the protein once it’s outside its natural cellular environment. The BCs, on the other hand, are an unstructured region that can freely move and that could resemble the flexibility properties of the prenyl group more closely than other folded membrane binding domains do. (2) The BCs are mainly positively charged, resembling the C-terminal polybasic region (that is also associated with membrane-binding [Johnson et al., 2012]). Thereby the C-terminal addition of BCs to Cdc42 may simply extend its positively charged C-terminal region without introducing other properties to it.

We designed a Cdc42 construct (Cdc42-BC) similar to Meca *et al*. [Meca et al., 2019]), where two out of three BC regions (51 aa) got inserted into the protein’s C-terminus in between the polybasic region and the CAAX box [Tschirpke et al., 2023]. Cdc42-BC recombinantly expressed in similar yield as other Cdc42 constructs, could be purified in a one-step His-affinity chromatography, and showed a normal GTPase activity and interaction with the GEF Cdc24. A fluorescent version of Cdc42-BC could also be generated (here the fluorescent protein mNeongreen is appended to a solvent-exposed loop of Cdc42 [Bendezú et al., 2015]) [Tschirpke et al., 2023].

However, initial membrane-binding experiments using fluorescence microscopy and fluorescence recovery after photobleaching suggest that Cdc42-BC does not sufficiently bind to membranes minicing the composition of the yeast plasma membrane (75% phosphatidylcholine, 20% phosphatidylserine, 5% phosphatidylinositol 4,5-bisphosphate [Meca et al., 2019]) (Chapter 5 in [Tschirpke, 2022]). Further, liposome co-flotation assays showed that the majority of Cdc42-BC does not bind to membranes (S1 Fig. 7). In contrast, Cdc42 farnesylated in a Sortase-mediated reaction does bind to such membranes (Fig. 3).

**S1 Figure 7.**
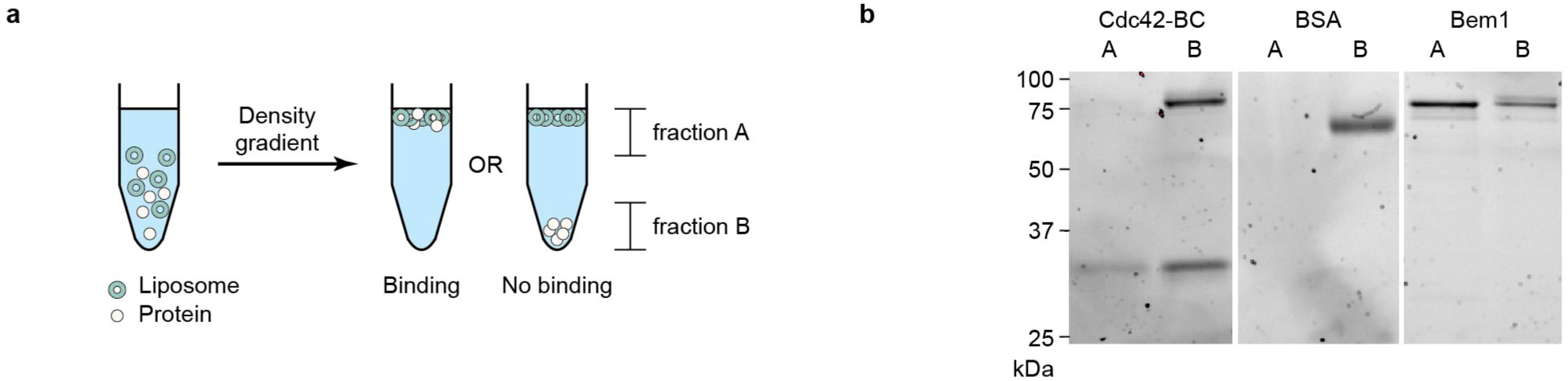
The majority of Cdc42-BC does not bind to membranes. (a) Illustration of liposome co-flotation assays. (b) SDS-PAGE analysis of the bound and unbound fraction of Cdc42-BC, Bem1 (positive control), and bovine serum albumin (BSA, negative control) in liposome co-flotation assays: Cdc42-BC is visible as two bands on SDS-PAGE; a lower (monomeric) and a higher (dimer) band [Tschirpke et al., 2023]. Only a small fraction of the monomeric Cdc42-BC band is visible in the bound fraction (fraction A), whereas the majority of protein remains in the unbound fraction (fraction B). In contrast, more than half of Bem1 is in the bound fraction. BSA does not bind to membranes. The co-flotation assay was carried out as described (materials and methods), with the following modifications: (1) The assays were carried out in absence of GTP, as the BC clusters (as well as Bem1) do not require GTP for membrane binding [Meca et al., 2019]. (2) Because no detergent was present in the buffers, no Bio-beads were added. (3) The membrane composition was adjusted to include phosphatidylinositol 4,5-bisphosphate, which was shown to be important for BC mediated membrane binding [Meca et al., 2019].

## Supplement S2

**S2 Figure 1.**
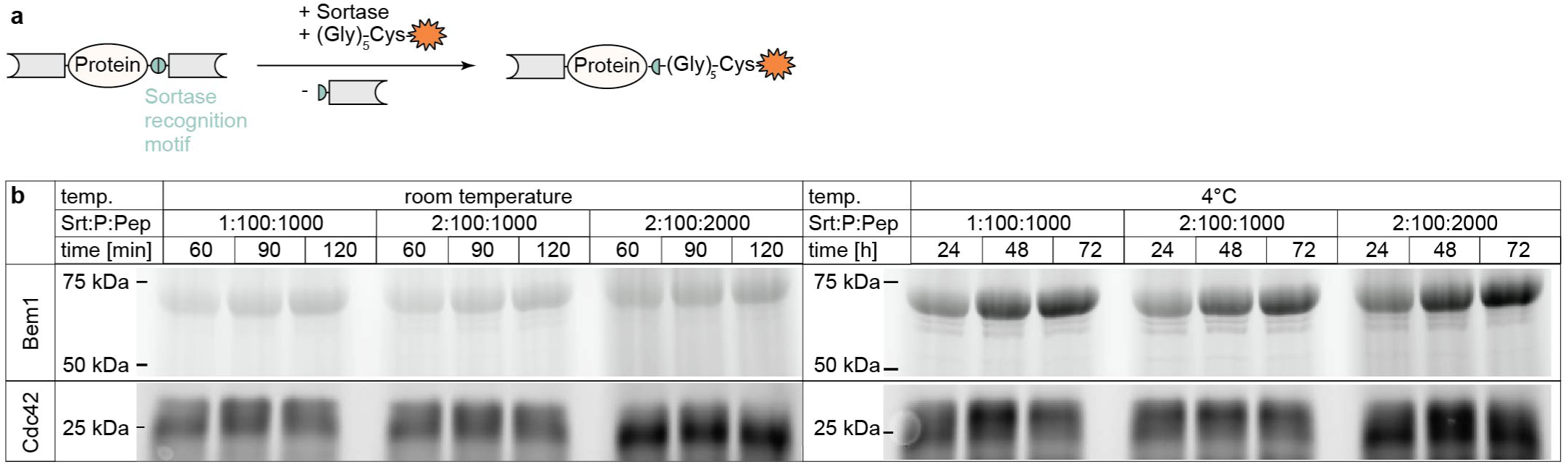
Condition screen for Sortase-mediated labelling of Cdc42 and Bem1 using Alexa488 peptide. (a) Schematic illustration of the Sortase-mediated labelling reaction of double-tagged protein (Cdc42 or Bem1) with Alexa488 peptide. (b) Condition screen of the labelling reaction of Bem1 (70 kDa) or Cdc42 (29 kDa) with Alexa488 peptide, using stated temperatures, Sortase : protein : Alexa 488 peptide (Srt:P:Pep) ratios, and incubation times. SDS-PAGE of the Alexa488-signal are shown.

## Supplement S3

**S3 Figure 1.**
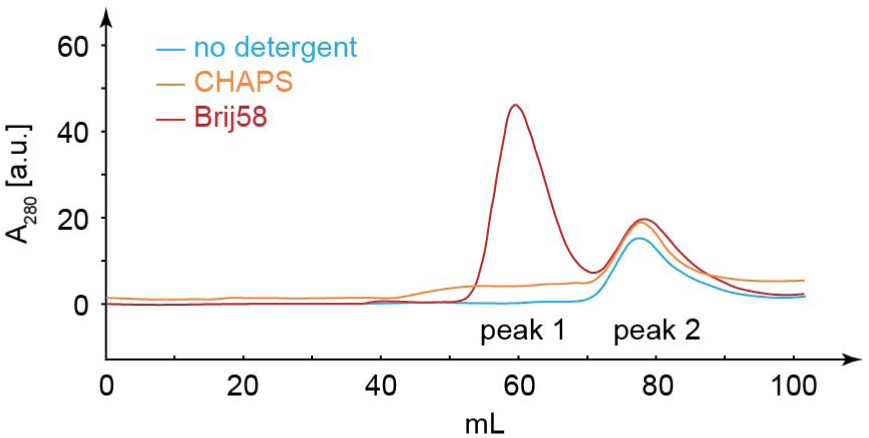
Addition of the detergent Brij58 results in an additional peak during SEC of Cdc42 samples after Sortase-mediated farnesylation. The labelling buffer / SEC buffer were supplemented with no detergent (blue), 2% / 0.5% CHAPS (orange), or 1% / 0.1% Brij58 (red).

## Supplement S4

His-AC flow-through samples of both SEC peak 1 and SEC peak 2 (Fig. 2) were dialyzed in (a) SEC buffer (Tab. 2) supplemented with 0.1% *n*-Dodecyl *β*-D-maltoside (DDM), (b) SEC buffer supplemented with 0.1% CHAPS, and (c) SEC buffer. Samples were sent to Max Perutz Labs Vienna for mass analysis. Independent of the detergent used, Flag-Cdc42-farnesyl (with a cleaved methionine: 28364.95 Da) was detected in all samples (S4 Tab. 1).

**S4 Table 1.** Mass analysis of His-AC flow-through (FT) of SEC peak 1 and SEC peak 2. Flag-Cdc42-farnesyl, with a cleaved methionine (28364.95 Da), is used as reference to calculate the error.

| Sample | Buffer | Detected mass [Da] | Error [Da] | Data shown in |
| --- | --- | --- | --- | --- |
| His-AC FT SEC peak 1 | SEC buffer + DDM | 28364.2 | -0.75 | S4 Fig. 1a |
| His-AC FT SEC peak 1 | SEC buffer + CHAPS | 28364.0 | -0.95 | S4 Fig. 1b |
| His-AC FT SEC peak 1 | SEC buffer | 28363.9 | -1.05 | S4 Fig. 1c |
| His-AC FT SEC peak 2 | SEC buffer + DDM | 28366.6 | 1.65 | S4 Fig. 2a |
| His-AC FT SEC peak 2 | SEC buffer + CHAPS | 28366.3 | 1.35 | S4 Fig. 2b |
| His-AC FT SEC peak 2 | SEC buffer | 28366.6 | 1.65 | S4 Fig. 2c |

**S4 Figure 1.**
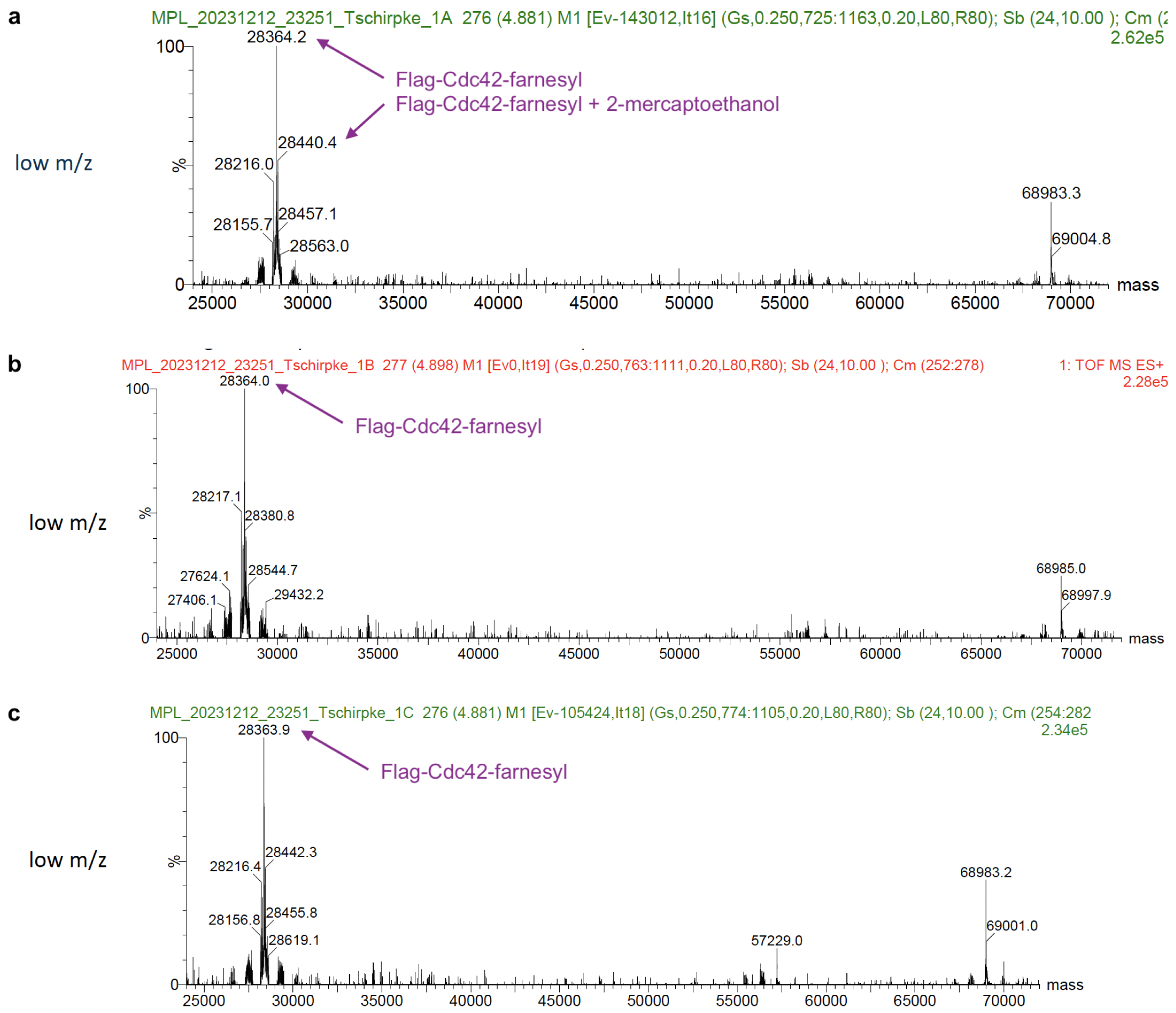
His-AC flow-through samples of SEC peak 1 contains Flag-Cdc42-farnesyl. Deconvoluted mass analysis spectrum of His-AC flow-through of SEC peak 1, dialyzed in (a) SEC buffer supplemented with 0.1% DDM, and (b) SEC buffer supplemented with 0.1% CHAPS, (c) SEC buffer. Mass analysis was conducted by Max Perutz Labs Vienna.

**S4 Figure 2.**
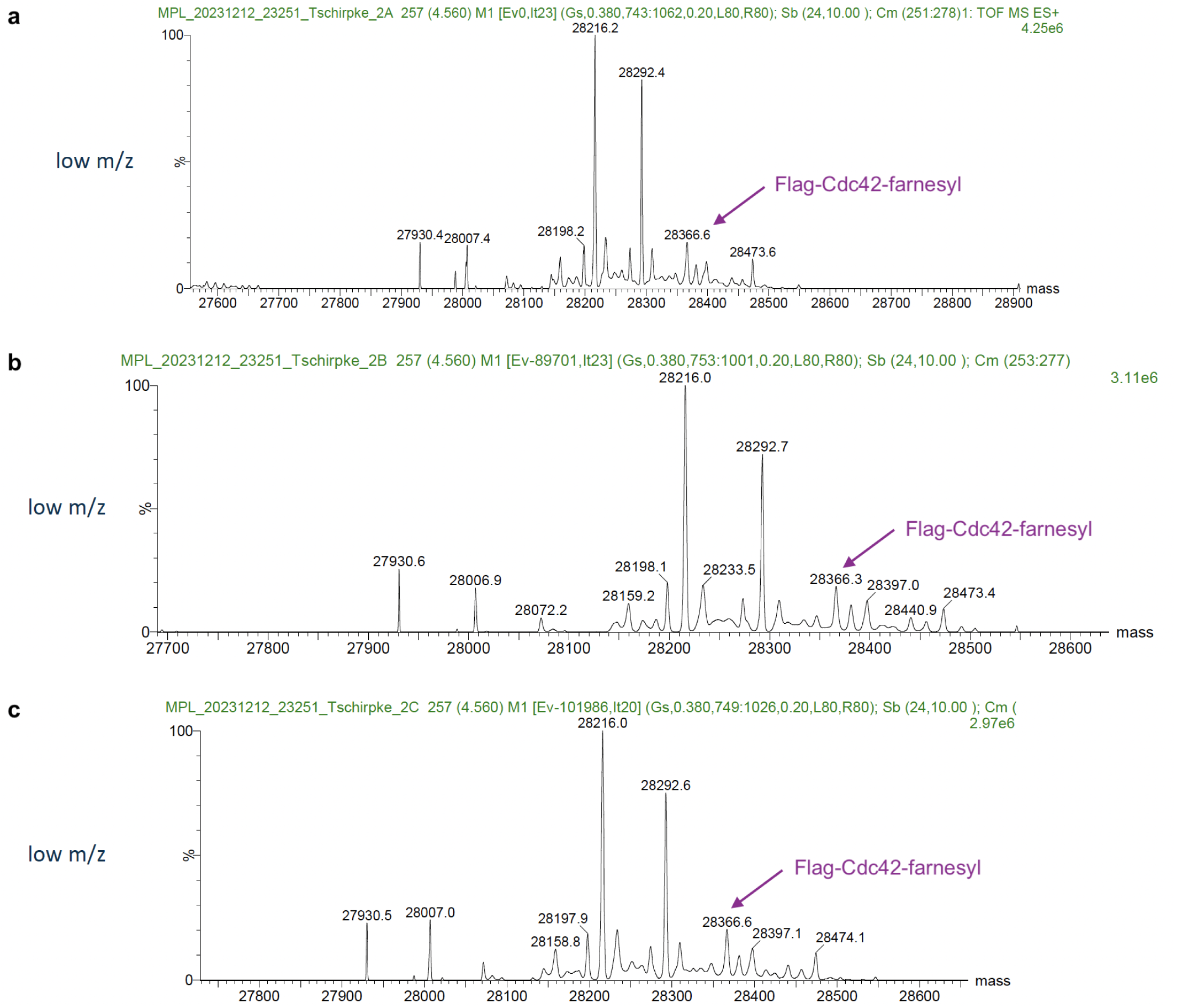
His-AC flow-through samples of SEC peak 2 contains Flag-Cdc42-farnesyl. Deconvoluted mass analysis spectrum of His-AC flow-through of SEC peak 2, dialyzed in (a) SEC buffer supplemented with 0.1% DDM, and (b) SEC buffer supplemented with 0.1% CHAPS, (c) SEC buffer. Mass analysis was conducted by Max Perutz Labs Vienna.

## Supplement S5

Cdc42 with an N-terminal 6His-tag and C-terminal Flag-tag (His-Cdc42-Flag) was labeled with farnesylated triglycine, using molar reaction ratios of Sortase : Cdc42 : farnesylated triglycine of 2:100:2’000 (S5 Fig. 1a). During the labeling reaction the Sortase enzyme can cleave Cdc42’s C-terminal tag and ligate farnesylated triglycine to the protein. The final reaction mixture therefore contains three Cdc42 species:

- educt: His-Cdc42-Flag, 29 kDa (marked with a diamond in S5 Fig. 1)
- desired product: His-Cdc42-farnesyl, 26.5 kDa (marked with a red dot in S5 Fig. 1)
- undesired side-product: His-Cdc42, 26 kDa (marked with a filled diamond in S5 Fig. 1)

Additionally, Sortase (22 kDa, but with an apparent size of 26 kDa (Fig. 2d)), cleaved C-terminal Flag-tags, and remaining farnesylated triglycine are part of the reaction mixture (S5 Fig. 1a,b).

As a first step, size exclusion chromatography (SEC) was used to separate reactants and products by size. Considering the proteins’ sizes, SEC separates a mixture of the three Cdc42 species and Sortase enzyme from remaining peptide and cleaved Flag-tag peptides (S5 Fig. 1b).

The protein-containing SEC peak was loaded onto a hydrophobic interaction chromatrography (HIC) column. Farnesyl is a very hydrophobic molecule, therefore farnesylated Cdc42 should bind way stronger to the hydrophobic column material in comparison to not farnesylated Cdc42 and Sortase. The elution profile showed two strongly overlapping peaks, indicating the presence of two distinct species (S5 Fig. 1b). To test if Cdc42 by itself can bind to the HIC column, we ran a HIC purification protocol using only a small amount of Cdc42. Cdc42 eluted in one broad peak that mostly overlaped with the second HIC peak (S5 Fig. 1b), suggesting that the second HIC peak (fractions C-E) contains the majority of educt and undesired product. We analyzed the reaction and purification steps using SDS-PAGE and anti-His and anti-Flag Western blotting (S5 Fig. 1c): Bands that are visible on the anti-His blot, but not on the anti-Flag blot, correspond to the undesired side-product (His-Cdc42, filled diamond). Bands that are visible on both anti-His and anti-Flag blots correspond to the educt (His-Cdc42-Flag, diamond). Our previous analysis (Fig. 2c,d) showed that farnesylated Cdc42 (red dot) runs at the same position as Cdc42 educt. Therefore, we are looking for a band on SDS-PAGE that runs at the same height as the Cdc42 educt, but that is not visible on the anti-Flag blot.

After 72 hours of reaction, we observe three bands on SDS-PAGE (S5 Fig. 1c): 25, 23, and 20 kDa, all of which were smaller than expected. It is possible that buffer components caused the samples to migrate lower than anticipated. The SEC peak contains two bands — 25 and 20 kDa — likely because the 23 and 20 kDa bands merged during this step. Western blotting shows that the 20 kDa band has a His-tag, while the 25 kDa band contains both His- and Flag-tags. This suggests the 20 kDa band corresponds to His-Cdc42, and the 25 kDa band is His-Cdc42-Flag. The 20 kDa band is the dominant species, while the 25 kDa band is faint, indicating that the major reaction product is the side-product His-Cdc42.

The fractions of the two overlapping HIC elution peaks (S5 Fig. 1b) show only a 20 kDa band on SDS-PAGE (S5 Fig. 1c), with increasing intensity in later elution volumes (from fraction A to E), supporting our hypothesis that the second HIC peak represents not farnesylated Cdc42. Both anti-His and anti-Flag Western blots show a similar increase in intensity from fractions A to E. The 20 kDa band visible on SDS-PAGE is detected only by the anti-His blot, confirming it is His-Cdc42. However, an additional, but weak, 25 kDa band appears on the anti-Flag blot, with its intensity also increasing from fraction A to E. The 25 kDa band, which corresponds to His-Cdc42-Flag, was likely present in such small amounts that it was not visible on SDS-PAGE and only detectable via Western blotting. This suggests that the second HIC peak contains both His-Cdc42 and His-Cdc42-Flag. The first HIC peak contains only minor amounts of His-Cdc42 and His-Cdc42-Flag, which are insufficient to explain the absorbance / peak height observed during HIC. This suggests that the first HIC peak likely includes farnesylated Cdc42, along with small quantities of His-Cdc42 and His-Cdc42-Flag. The presence of farnesylated Cdc42 in the first HIC peak is surprising, as we expected farnesylated Cdc42 to bind stronger to the column and thus elute in the second (not first) HIC peak. Overall, the significant overlap between the peaks points to poor separation of farnesylated and not farnesylated Cdc42 species. Our results partially align with those of Fres *et al*. [Fres et al., 2010], who found that different Butyl Sepharose column variants had varying abilities to separate farnesylated from not farnesylated protein.

**S5 Figure 1.**
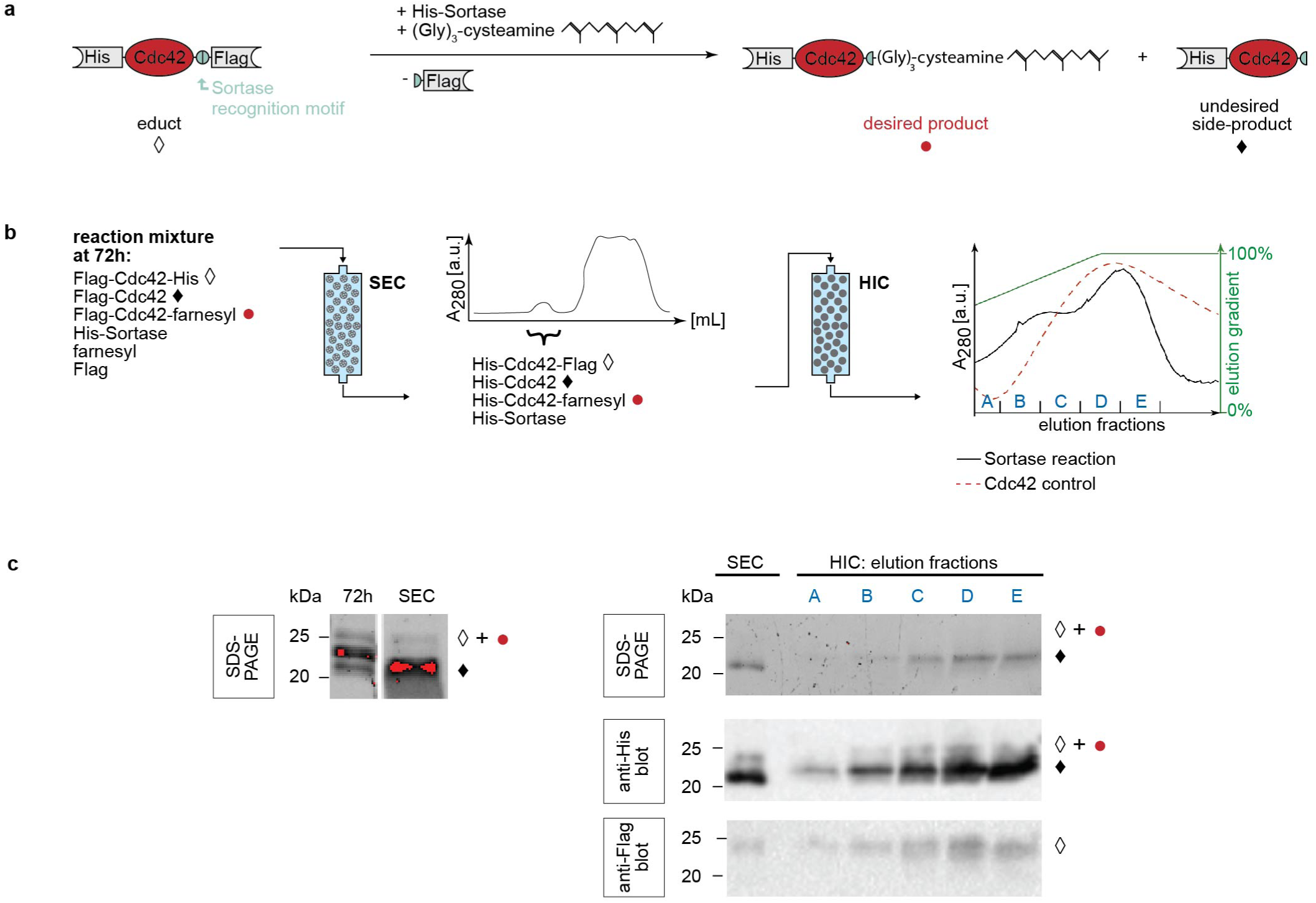
SEC and HIC lead to poor separation of farnesylated and not farnesylated Cdc42 species from Sortase-mediated farnesylation reactions. (a) Schematic illustration of the Sortase-mediated labeling reaction of double-tagged Cdc42 (His-Cdc42-Flag, diamond) with farnesylated triglycine: in addition to farnesylated Cdc42 (red dot), Cdc42 with a cleaved Flag-tag (filled diamond) can also occur as a side-product. (b) Clean-up procedure for the labeling reaction of His-Cdc42-Flag with farnesylated triglycine (farnesyl): after an incubation of 72 h size exclusion chromatography (SEC) is used to separate unreacted farnesylated triglycine and cleaved Flag-tags from proteins, which elute in one peak. This SEC peak is loaded onto a hydrophobic interaction chromatography (HIC) column. The HIC elution profile of SEC peak (black line, fractions A-E) overlaps with the elution profile of a pure His-Cdc42-Flag control sample (dashed red line), suggesting that HIC is not a suitable method to separate farnesylated Cdc42 from unreacted Cdc42. (c) SDS-PAGE, anti-His and anti-Flag Western blot of the reaction mixture post-reaction (72 h), the SEC peak, and HIC fractions A-E.

## Supplement S6

**S6 Figure 1.**
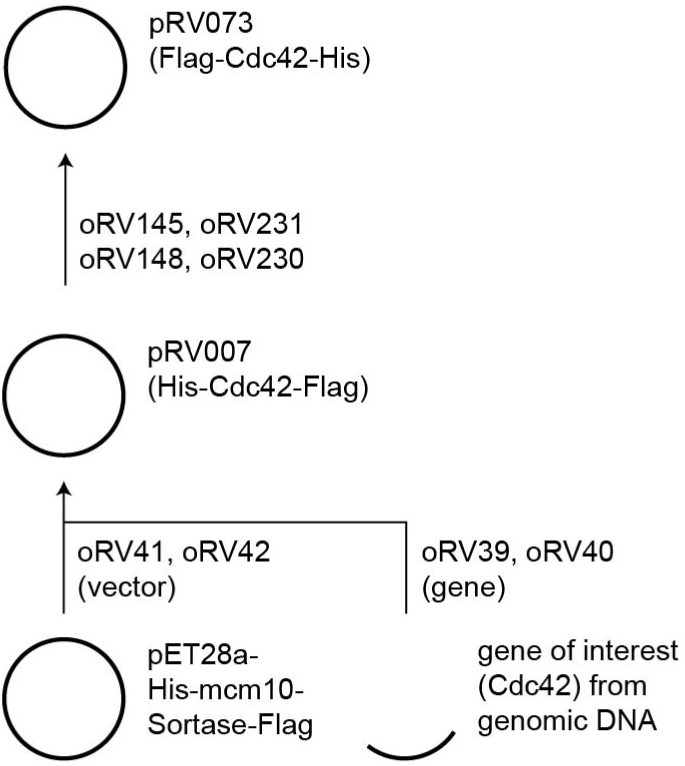
Schematic of the plasmid construction. pET28a-His-mcm10-Sortase-Flag was received from N. Dekker (TU Delft) and is based on pBP6 [Douglas and Diffley, 2016].

**S6 Table 1.**
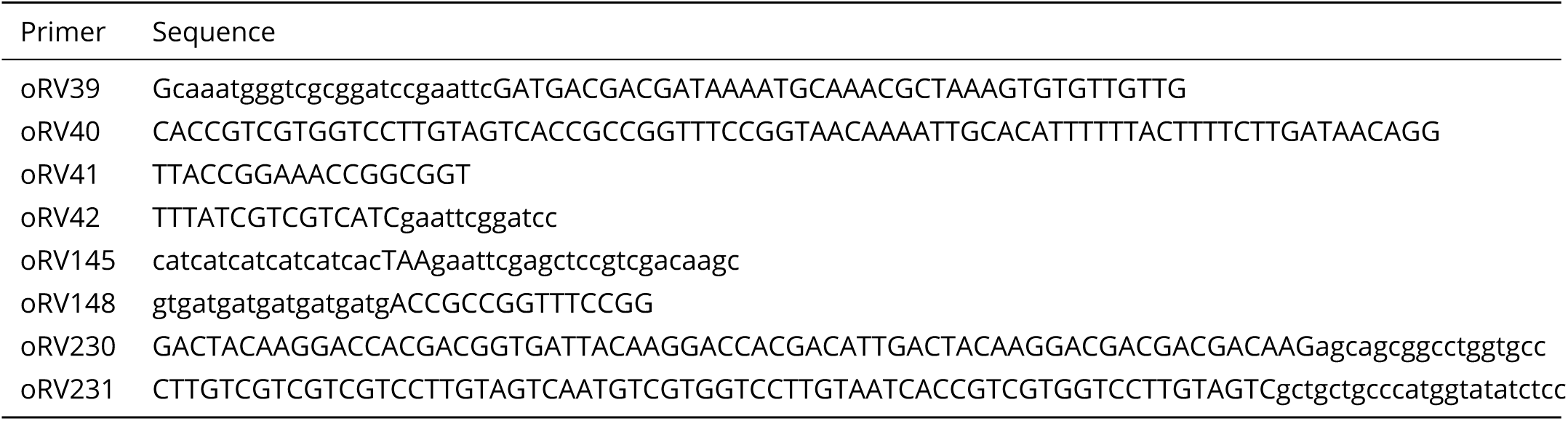
Primer overview.

**Protein sequences:**

His-Cdc42-Flag, pRV007:

MGSSHHHHHHSSGLVPRGSH<u>MASMTGGQQMGRGSEF</u>DDDDKMQTLKCVVV GDGAVGKTCLLISYTTNQFPADYVPTVFDNYAVTVMIGDEPYTLGLFDTA GQEDYDRLRPLSYPSTDVFLVCFSVISPPSFENVKEKWFPEVHHHCPGVP CLVVGTQIDLRDDKVIIEKLQRQRLRPITSEQGSRLARELKAVKYVECSA LTQRGLKNVFDEAIVAALEPPVIKKSKKCAILLPETGGDYKDHDGDYKDH DIDYKDDDDK

Flag-Cdc42-His, pRV073:

MGSSDYKDHDGDYKDHDIDYKDDDDKSSGLVPRGS<u>HMASMTGGQQMGRGS</u> <u>EF</u>DDDDKMQTLKCVVVGDGAVGKTCLLISYTTNQFPADYVPTVFDNYAVT VMIGDEPYTLGLFDTAGQEDYDRLRPLSYPSTDVFLVCFSVISPPSFENV KEKWFPEVHHHCPGVPCLVVGTQIDLRDDKVIIEKLQRQRLRPITSEQGS RLARELKAVKYVECSALTQRGLKNVFDEAIVAALEPPVIKKSKKCAILLP ETGGHHHHHH

The protein constructs contain the following features:

- 6His-tag: HHHHHH
- Flag-tag: DYKDHDGDYKDHDIDYKDDDDK
- Thrombin cut site: LVPRGS
- Enterokinase cut site: DDDDK
- Sortase recognition motif: LPETG
- T7 tag (to aid protein expression): MASMTGGQQMGRGSEF

More information on the protein constructs is given in [Tschirpke et al., 2023].

**S6 Figure 2.**
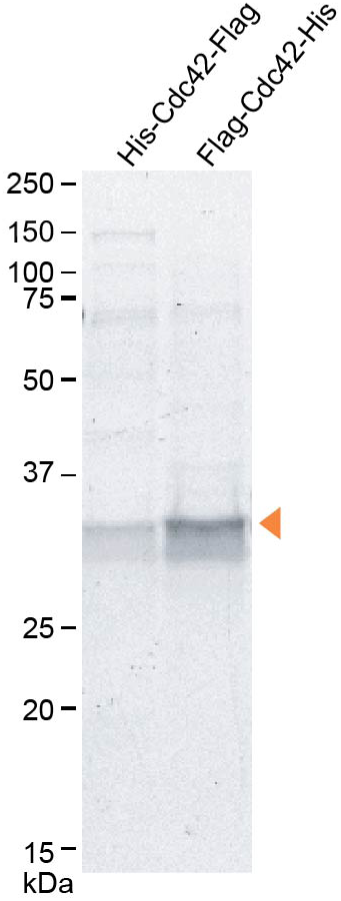
SDS-PAGE of used proteins. An orange arrow indicates the band of the correct size.

## Supplement S7

**S7 Figure 1.**
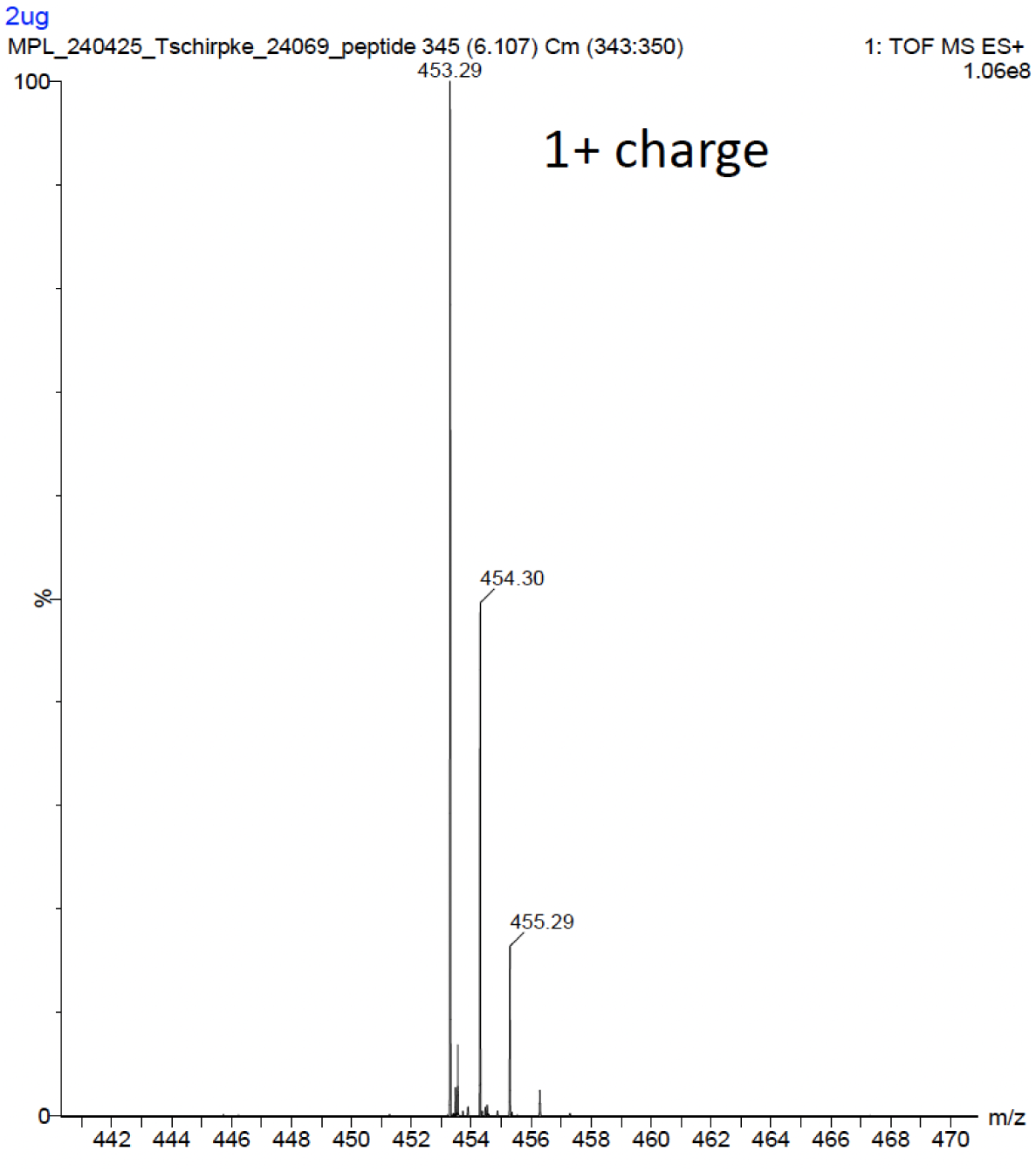
Deconvoluted mass analysis spectrum of H_2_N-Gly_3_-cysteamine-farnesyl (farnesyl peptide), obtained by Max Perutz Labs Vienna.

## Footnotes

1 By the time we started carrying out experiments peptides with farnesyl modification were commercially not available yet, though several companies offer them by now.

2 We also detected some of the 30 kDa band (educt Flag-Cdc42-His) in the His-AC flow-through, likely due to incomplete binding to the column.

3 Initially we intended to monitor Cdc42 farnesylation using anti-farnesyl Western blotting, but an eight condition screen (combination of two primary anti-bodies, two secondary antibodies, and two detection kit solutions that were successfully used to detect farnesyl previously [Kennedy et al., 2019, Li et al., 2020]) revealed that all conditions showed false-positive and false-negative bands for farnesyl (data not shown).

4 The bands at 50 kDa correspond to Cdc42 dimers that form in denaturing conditions [Tschirpke et al., 2023].

5 The different expression conditions can not be directly compared to each other, as the images are from distinct Western blots. To compare the expression levels of the three conditions, see S1 Fig. 6b. Here three conditions are shown in the same blot (for His-Cdc42:CTIS).

